# Temporal and genomic analysis of additive genetic variance in breeding programmes

**DOI:** 10.1101/2020.08.29.273250

**Authors:** Letícia A. de C. Lara, Ivan Pocrnic, R. Chris Gaynor, Gregor Gorjanc

## Abstract

This study demonstrates a framework for temporal and genomic analysis of additive genetic variance in a breeding programme. Traditionally we used specific experimental designs to estimate genetic variance for a specific group of individuals and a general pedigree-based model to estimate genetic variance for pedigree founders. However, with the pedigree-based model we can also analyse temporal changes in genetic variance by summarising sampled realisations of genetic values from a fitted model. Here we extend this analysis to a marker-based model and build a framework for temporal and genomic analyses of genetic variance. The framework involves three steps: (i) fitting a marker-based model to data, (ii) sampling realisations of marker effects from the fitted model and for each sample calculating realisations of genetic values, and (iii) calculating variance of the sampled genetic values by time and genome partitions. Genome partitions enable estimation of contributions from chromosomes and chromosome pairs and genic and linkage-disequilibrium variances. We demonstrate the framework by analysing data from a simulated breeding programme involving a complex trait with additive gene action. We use the full Bayesian and empirical Bayesian approaches to account for the uncertainty due to model fitting. We also evaluate the use of principal component approximation. Results show good concordance between the simulated and estimated variances for temporal and genomic analyses and give insight into genetic processes. For example, we observe reduction of genic variance due to selection and drift and buildup of negative linkage-disequilibrium (the Bulmer effect) due to directional selection. In this study the popular empirical Bayesian approach estimated the variances well but it underestimated uncertainty of the estimates. The principal components approximation biases estimates, in particular for the genic variance. This study gives breeders a framework to analyse genetic variance and its components in different stages of a programme and over time.

## 1 Introduction

In this study we analyse temporal and genomic trends of additive genetic variance in different stages of a breeding programme. Genetic variance is one of the critical parameters in a breeding programme because it determines the potential for selection (Lush, 1937; Falconer and Mackay, 1996; Lynch and Walsh, 1998; Walsh and Lynch, 2018). Estimation of genetic variance has therefore received considerable attention in the literature (Lynch and Walsh, 1998; Walsh and Lynch, 2018). Most of the attention in literature is on statistical models and approaches for estimation. Surprisingly, far less attention has been given to temporal trends in genetic variance, even though such trends indicate sustainability of a breeding programme. Recent ability to observe genomes at scale has renewed interest in analysing genetic variance. In this study we show that with a combination of established and new approaches we can use a simple framework to analyse temporal and genomic trends in genetic variance in a breeding programme.

Estimation of genetic variance in breeding programmes has a long history and a recent revival with the advent of genomic information. Historically, genetic variance was estimated with an analysis of variance (ANOVA) methods in tailored experimental designs ranging from simple parent-offspring or sib groups to North Carolina and diallel designs (Falconer and Mackay, 1996; Lynch and Walsh, 1998; Bernardo, 2002; Awata *et al*., 2018). With these designs we partition phenotypic variance into variance between and within groups and “translate” these components into genetic variance based on expected genetic relationships within and between groups. Animal breeders have soon moved from experimental designs to a general pedigree-based model to analyse their observational data (Henderson, 1976). Plant breeders generally analyse experimental data and have only recently started to adopt the pedigree-based model (Oakey *et al*., 2006, 2007; Piepho *et al*., 2008). There are many logistical and conceptual reasons for this. One reason is that with the pedigree-based model we estimate genetic variance between the founders of a pedigree (Sorensen and Kennedy, 1984; Kennedy *et al*., 1988), while genetic variance between their descendants is arguably more relevant for breeding (Piepho *et al*., 2008). The advent of genomic information revived interest in the estimation of genetic variance and spurred active development of genome-based models (Bernardo, 1994, 1996; Meuwissen *et al*., 2001; VanRaden, 2008). The genome-based model replaces expected relationships from the experimental designs or pedigree with realised relationships measured by marker genotypes. The estimate of genetic variance from the genome-based model pertains to all genotyped individuals (Hayes *et al*., 2009) and can be obtained using either a genome-based model with genetic values or a genome-based model with marker effects (marker-based model) (Strandén and Garrick, 2009). We note though that the resulting “genomic variance” is at odds with the quantitative genetics definition of genetic variance (Gianola *et al*., 2009; de los Campos *et al*., 2015). Specifically, the genome-based model is defined with the (scaled) variance of marker effects and not with genetic variance. Further, markers are not necessarily quantitative trait loci affecting phenotype. Both of these points lead to model “misspecification” in a sense that model parameters do not represent quantitative genetic parameters (Gianola *et al*., 2009; de los Campos *et al*., 2015). We will come back to this note repeatedly.

In parallel to the development of data sources and corresponding statistical models, there has been active development in statistical and computational approaches for the estimation of genetic variance. The three most used are method of moments, likelihood and Bayesian approach. The method of moments that is used with the ANOVA is computationally simple but can yield biased estimates outside of the parameter space. It also does not generalise to unbalanced data. The likelihood approach has better statistical properties than the method of moments (Sorensen and Gianola, 2007). With the likelihood approach we specify a probability distribution for observed data and find the most likely value of model parameters that would give rise to the observed data. Use of this approach to estimate genetic variances is extensively described in Meyer (1985); Meyer and Hill (1997); Smith *et al*. (2005); Thompson *et al*. (2005); Thompson (2019). The Bayesian approach improves the likelihood approach in two ways. First, it incorporates prior knowledge (distribution) for all model parameters (means and variances), which can improve estimation (Sorensen and Gianola, 2007; Hem *et al*., 2020). Second, it treats all model parameters in a probabilistically consistent manner such that estimation uncertainty is propagated to all estimated model parameters (Sorensen and Gianola, 2007). The full probabilistic treatment makes the Bayesian approach computationally more demanding than the likelihood approach. We commonly handle the computational demand by using an empirical Bayesian approach where we first estimate most likely values for variance parameters and conditional on these estimate other model parameters (Efron, 1996; Sorensen and Gianola, 2007). In the marker-based model, the empirical Bayesian approach estimates model variances from the data at hand and conditional on these estimates all marker effects jointly to account for uncertainty of estimating marker effects (uncertainty of estimating model variances is ignored). The full Bayesian approach accounts for uncertainty in estimating model variances and marker effects. The full Bayesian approach is commonly approached with computationally intensive sampling methods such as Monte Carlo Markov Chain (MCMC) (Gilks *et al*., 1995; Brooks *et al*., 2011). MCMC on genome-based models with many individuals or markers can be time-consuming. To this end various dimensionality-reduction approaches have been proposed, for example, singular value decomposition (SVD) of marker genotypes where we fit a small number of principal components that capture majority of variance in marker genotypes (Tusell *et al*., 2013; Ødegård *et al*., 2018).

Variances from pedigree and genome-based models do not inform about temporal and genomic trends in genetic variance because they pertain to a specific group of individuals and encompass the whole genome (Sorensen and Kennedy, 1984; Kennedy *et al*., 1988; Hayes *et al*., 2009). However, these models can be used for temporal and genomic analyses of genetic variance with some post-processing. Sorensen *et al*. (2001) showed how to analyse the temporal trend in genetic variance. They fitted a pedigree-based model and inferred genetic variance for several time partitions by sampling realisations of genetic values from the fitted model and calculating variance of the realisations partitioned in time groups. They used the Bayesian approach and MCMC, but their concept is general and can be used with other statistical and computational approaches. The important distinction here is between model fitting to estimate statistical/model parameters and post-processing to estimate quantitative genetics parameters. This distinction enables flexibility to fit a generic model, for example LASSO (Tibshirani, 1996), and to estimate quantitative genetics parameters from post-processing results of the model. This gives a potential to (partially) address the issue of “misspecification” with genome-based models (Gianola *et al*., 2009; de los Campos *et al*., 2015). Partially, because we need enough markers to capture all variation at quantitative trait loci. Lehermeier *et al*. (2017) used the same approach with the marker-based model and analysed the contribution of linkage-disequilibrium to genetic variance. Recently, Allier *et al*. (2019) also used the marker-based model on data from a maize breeding programme to infer trends in genetic mean and genetic variance as well as the contribution of allele diversity (genic variance) and of linkage-disequilibrium to genetic variance (Bulmer, 1971; Lynch and Walsh, 1998; Walsh and Lynch, 2018).

The aim of this work is to i) build and validate a flexible framework based on the work of Sorensen *et al*. (2001), Lehermeier *et al*. (2017) and Allier *et al*. (2019), ii) show how to evaluate temporal and genomic analysis of additive genetic variance in different stages of a breeding programme and iii) indicate genetic processes that change genome. We also show how different statistical approaches affect the results. To this end we have validated our work with a simulated breeding programme, used a marker-based model to estimate marker effects and based on these estimated temporal and genomic trends in additive genetic variance. The results show good concordance between the simulated and estimated variances and give insight into genetic processes. In this study the popular empirical Bayesian approach estimated variances well but it underestimated uncertainty of the estimates. The principal components approximation biased estimates, in particular for the genic variance.

## 2 Materials and Methods

In this section we present study material and methods in five parts: (1) simulation of a breeding programme where we generate true values and observed data, (2) temporal and genomic analysis of genetic variance where we demonstrate the framework assuming we know the true quantitative trait locus genotypes and their effects, (3) statistical analysis of observed data where we describe marker-based model fitted to observed data, (4) statistical and computational approaches to estimate marker effects, genetic values and variances, and (5) software implementation.

### 2.1 Breeding programme simulation

We simulated an entire wheat breeding programme considering additive genetic architecture for a quantitative trait. We have performed one simulation replicate for most analyses to focus on one dataset, but we also evaluated consistency of estimates for a subset of analyses on 10 simulation replicates. We followed a breeding programme described by Gaynor *et al*. (2017) with 21 years of a conventional phenotypic selection for yield (Fig. 1). We started with the simulation of whole-genome sequences for 21 chromosome pairs and extracted random 600 biallelic single nucleotide polymorphisms (SNP) as markers per chromosome and random 100 SNP as quantitative trait loci (QTL) per chromosome. We assumed that the 2,100 QTL had an additive effect on yield and sampled their effects from a normal distribution. We coded genotypes as 0 for reference homozygote, 1 for heterozygote and 2 for alternative homozygote. From the simulated whole-genome sequences, we created 70 inbred lines and crossed them to generate 100 biparental populations. Each population had 100 F_1_ that had their genome doubled and planted in headrows (altogether 10,000). In the headrows we visually evaluated the lines (trait heritability of 0.1) and advanced the best 500 into a preliminary yield trial. In the preliminary yield trial we evaluated the lines in an unreplicated trial (trait heritability of 0.2) and advanced the best 50 into an advanced yield trial. In the advanced yield trial we evaluated the lines in a small multi-location replicated trial (trait heritability of 0.5) and advanced the best 10 into an elite yield trial. In the elite yield trial we evaluated the lines for two consecutive years in a large multi-location replicated trial (trait heritability of 0.67) and released one variety. We used the best lines from the advanced and elite yield trials as parents to start a new breeding cycle.

**Figure 1:**
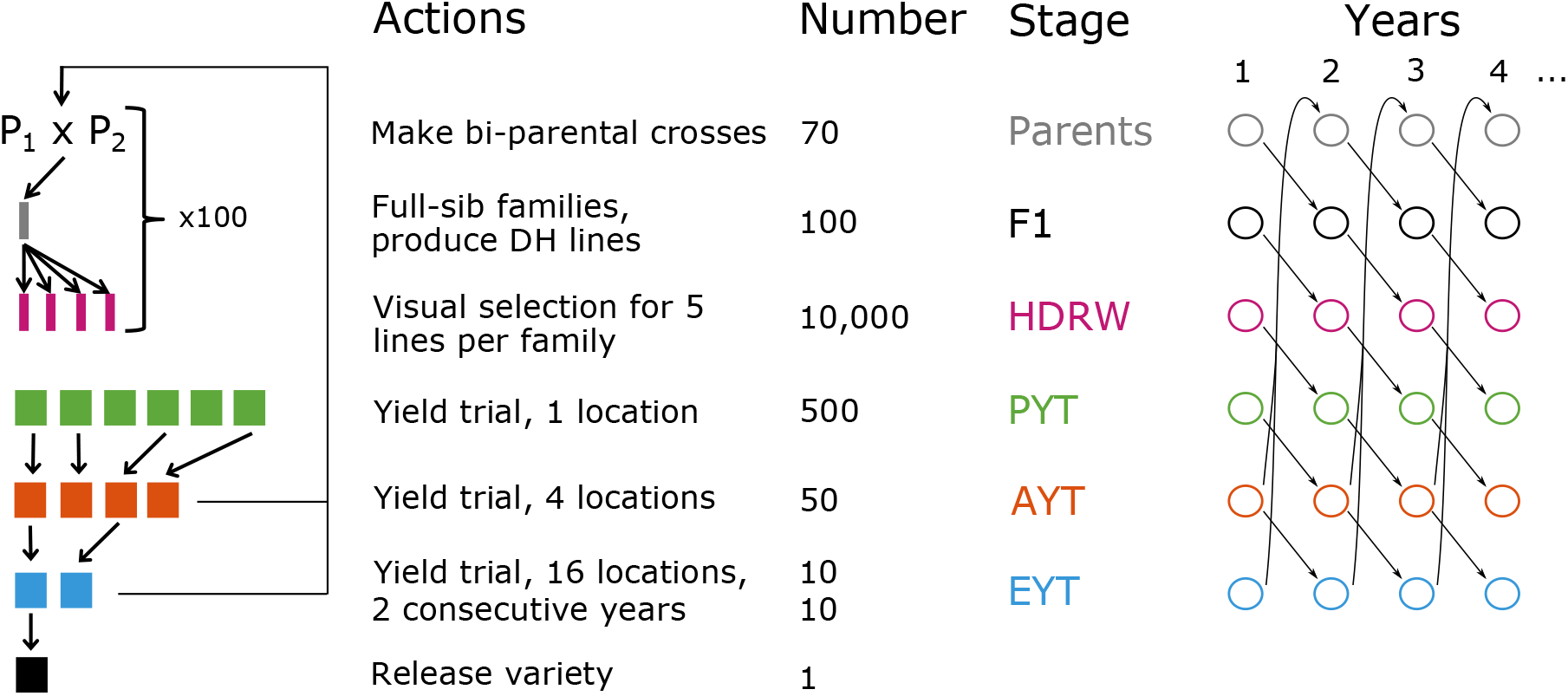
Simulated wheat breeding programme with parents, F_1_ progeny (F1), headrows (HDRW), preliminary yield trial (PYT), advanced yield trial (AYT), elite yield trial (EYT) and a released variety

Throughout the simulation we have saved phenotype and marker genotype data to generate a training population for genomic modelling. We did not use the genomic data in the simulation of a breeding programme, but only saved it for the statistical analysis of temporal and genomic trends of genetic variance. To this end, we have constructed a training population that spanned the last 6 years of the simulation, from year 16 to 21. This training population covered 3,070 lines with preliminary, advanced and elite yield trial phenotypes (altogether 3,420 phenotypes) and corresponding 10,500 marker genotypes.

### 2.2 Temporal and genomic analysis of genetic variation

Here we describe a flexible framework for temporal and genomic analysis of genetic variation, assuming that we know the QTL genotypes and their effects. In the following sub-sections, we estimate the temporal and genomic trends from observed phenotypes and marker genotypes and compare them to true values. The framework consists of four steps. First, we define whole-genome genetic values from QTL genotypes and their effects. Second, we partition individuals and their genetic values by time to calculate genetic variances over these time partitions for temporal analysis. Third, we partition whole-genome genetic values into chromosome and locus genetic values to calculate genetic variances and covariances over these genomic partitions for genomic analysis. This calculation involves three “layers” of variances: (a) total (whole-genome) genetic variance, (b) chromosome variances alongside linkage-disequilibrium covariances between chromosomes, and (c) locus genic variances alongside locus linkage-disequilibrium covariances within chromosomes and locus linkage-disequilibrium covariances between chromosomes. Fourth, we combine temporal and genomic analyses.

First, let ***Q*** be *n*_*i*_ × *n*_*q*_ matrix of QTL genotypes for *n*_*i*_ individuals at *n*_*q*_ QTL and ***α*** be *n*_*q*_ × 1 vector of QTL additive effects. Whole-genome genetic values of *n*_*i*_ individuals are a linear combination of QTL genotypes and their effects, ***a*** = ***Qα***. Variance of these values is genetic variance, 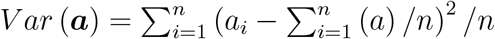. Note that this variance pertains to all *n*_*i*_ individuals and might not be an informative measure if these individuals span multiple stages and years of a breeding programme. In fact, any genetic trend or population structure will likely inflate this variance measure and mislead breeders in overestimating the amount of genetic variance. This is why we need temporal analysis of genetic variance.

Second, for the temporal analysis of genetic variance we partition the vector of genetic values by time and calculate variance for each time partition. For example, assume that individuals and their genetic values are ordered by time and that we partition them into time groups as ***a***[1 : *k*], ***a***[(*k* + 1) : *l*], ***a***[(*l* + 1) : *m*], … Then the temporal analysis of genetic variance is obtained by calculating variance for each time partition: 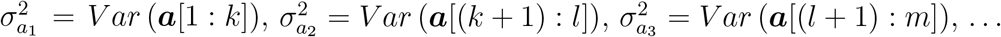

Third, for the genomic analysis of genetic variance we initially partition whole-genome genetic values ***a*** into an *n*_*i*_ × *n*_*c*_ matrix of *n*_*c*_ chromosome genetic values ***A***_*c*_ such that 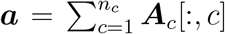. We obtain these chromosome genetic values by summing locus genetic values ***A***_*q*_ on each chromosome, ***A***_*c*_[*i, c*] = ∑_*l*_ ***Q***[*i, l*]***α***[*l*] for *l* running over 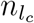 QTL on a chromosome *c*. Note that 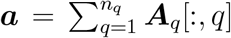 and 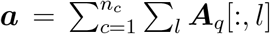 for *l* running over 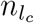 QTL on a chromosome *c*. To calculate genetic variances over these genomic partitions we will use the variance sum rule *Var*(*x* + *y*) = *Var*(*x*) + *Var*(*y*) + 2*Cov*(*x, y*) and the variance product rule *Var*(*xa*) = *Var*(*x*)*a*^2^. Partitioning of the genetic variance 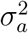 by chromosomes gives the sum of *n*_*c*_ chromosome variances 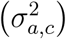 and *n*_*c*_ ∗ (*n*_*c*_ − 1) covariances between chromosomes (*σ*_(*a,c*′)(*a,c*)_):

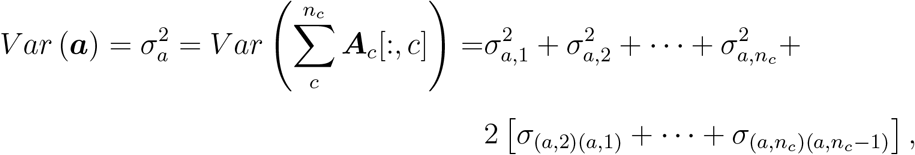

with covariances between chromosomes being between-chromosome linkage-disequilibrium covariances (Fig. 2). Partitioning of a chromosome genetic variance 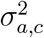 by loci gives the sum of 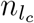 locus variances 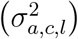 and *n*_*l*_ ∗ (*n*_*l*_ − 1) covariances between loci (*σ*_(*a,c,l*′)(*a,c,l*)_):

**Figure 2:**
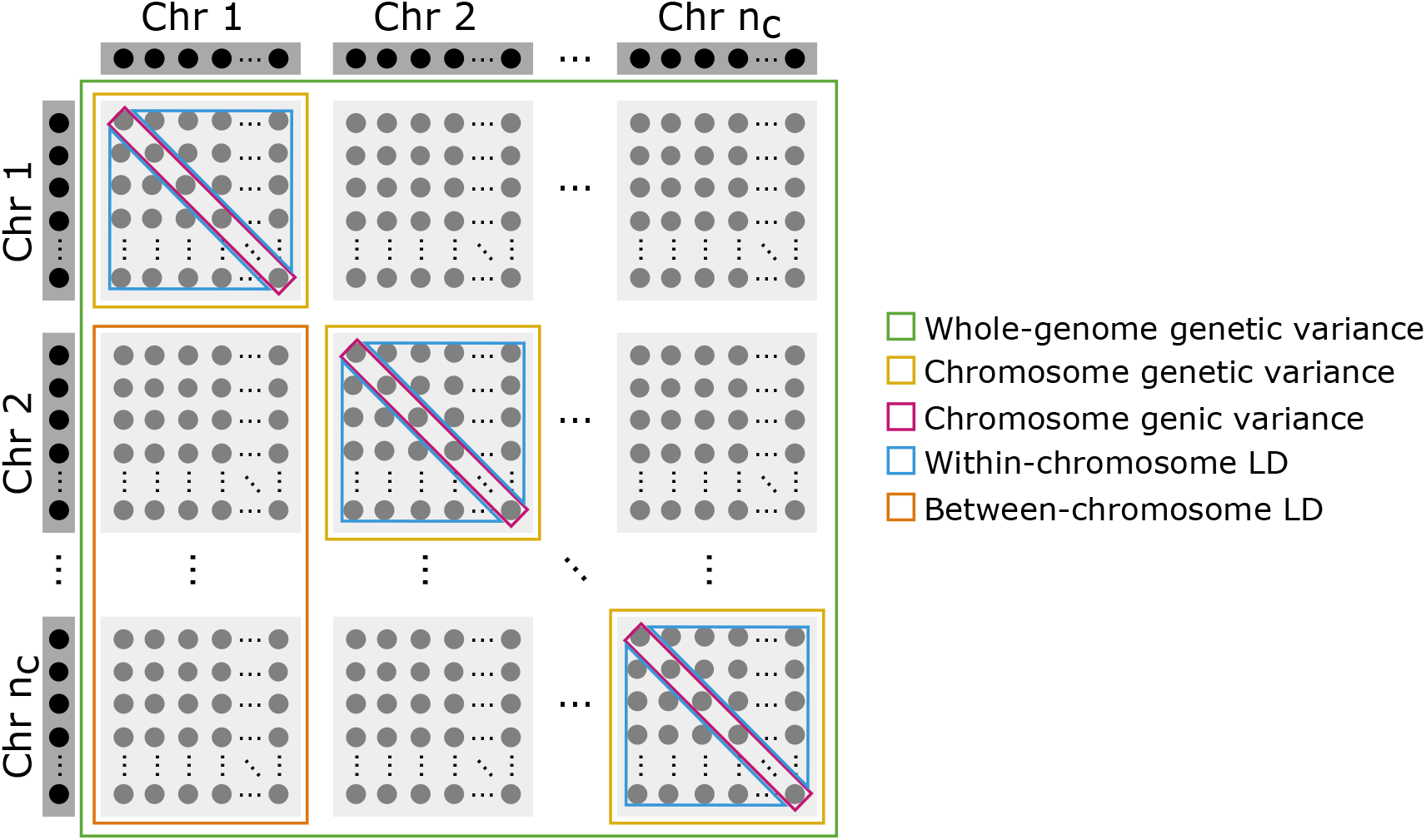
Illustrative scheme of genomic partitioning of whole-genome genetic variance by chromosomes and loci into genic, and within- and between-chromosome linkage-disequilibrium (LD) components

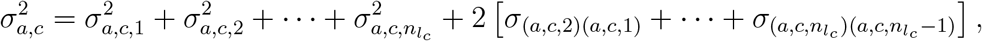

with locus variances being genic variances and covariances between loci being within-chromosome linkage-disequilibrium covariances (Fig. 2) (Bulmer, 1971; Lynch and Walsh, 1998; Walsh and Lynch, 2018). Locus genic variance is a function of variance in locus genotypes and their allele substitution effect (using variance product rule):

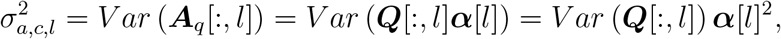

where we emphasise that we do not use the common Hardy-Weinberg assumption of *Var* (***Q***[:, *l*]) = 2*p*_*l*_(1−*p*_*l*_) with *p*_*l*_ being allele frequency. Instead, we advocate to calculate empirical variance in observed locus genotypes, *Var* (***Q***[:, *l*]). We will return to this point in discussion. Locus linkage-disequilibrium covariance is a function of covariance between genotypes at two loci and their allele substitution effects:

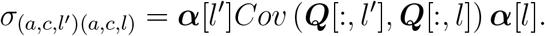

We can now partition the whole-genome genetic variance over chromosomes and loci as a sum of genic variances, within-chromosome linkage-disequilibrium covariances, and between-chromosome linkage-disequilibrium covariances (Fig. 2):

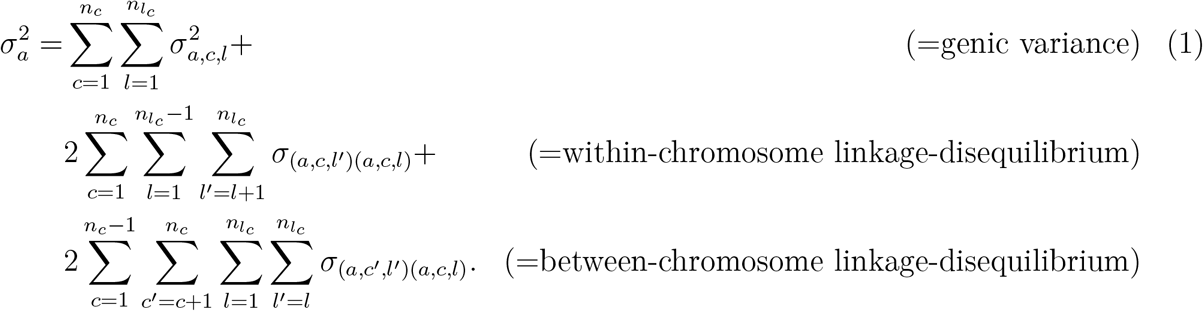

With *n*_*l*_ = 2, 100 QTL spread evenly over *n*_*c*_ = 21 chromosomes, the total number of locus combinations is *n*_*l*_ ∗ *n*_*l*_ = 4, 410, 000 and the total number of chromosome combinations is *n*_*c*_ ∗ *n*_*c*_ = 441. The framework partitions genetic variance into *n*_*l*_ = 2, 100 locus genic variances (*n*_*c*_ = 21 chromosome genic variances), 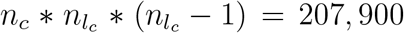 locus within-chromosome linkage-disequilibrium covariances (*n*_*c*_ = 21 chromosome within-chromosome linkage-disequilibrium covariances), and 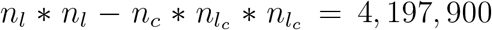 locus between-chromosome linkage-disequilibrium covariances (*n*_*c*_ ∗ *n*_*c*_ − *n*_*c*_ = 420 chromosome between-chromosome linkage-disequilibrium covariances). We emphasise these numbers because we often hear colleagues saying that there is no or limited between-chromosome linkage-disequilibrium (due to the lack of physical linkage). However, selection and other genetic processes generate within- and between-chromosome linkage-disequilibrium (Bulmer, 1971; Lynch and Walsh, 1998; Walsh and Lynch, 2018). Even if the between-chromosome linkage-disequilibrium covariances are small, there is a very large number of them and they can collectively have a sizeable effect on genetic variance as we show in results.

Fourth, for the joint temporal and genomic analysis, we perform genomic partitioning and variance calculations for individuals and their genetic values partitioned by time.

### 2.3 Statistical analysis of observed data

In the previous sub-section we assumed we know the QTL and their effects. In reality we observe phenotypes and marker genotypes and make inferences based on this information. To this end we fitted the marker-based model:

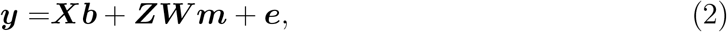

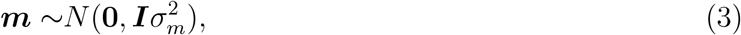

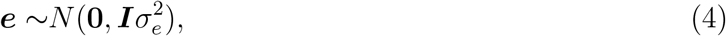

where, ***y*** is an *n*_*y*_ × 1 vector of *n*_*y*_ phenotypic values, ***X*** is an *n*_*y*_ × *n*_*b*_ incidence matrix for *n*_*b*_ intercept and year effects ***b, Z*** is an *n*_*y*_ × *n*_*i*_ incidence matrix for *n*_*i*_ lines whose marker genotype data is in an *n*_*i*_ × *n*_*m*_ matrix ***W*** for *n*_*m*_ marker effects ***m***, and ***e*** is an *n*_*y*_ × 1 vector of *n*_*y*_ residuals. In this study *n*_*y*_ was 3,420, *n*_*b*_ was 6, *n*_*i*_ was 3,070 and *n*_*m*_ was 10,500. We assumed that marker effects are *a priori* uncorrelated and normally distributed with zero mean and variance component describing variation between marker effects 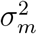 (Eq. 3). We further assumed that residuals are uncorrelated and normally distributed with zero mean and residual variance 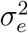 (Eq. 4). We ignored that different yield trials had different amount or replication and therefore different error variance.

The model (Eq. 2-4) has location parameters (means) ***b*** and ***m*** and dispersion parameters (variances) 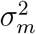 and 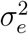. We emphasise that 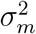 is variance between marker effects and note that the commonly used approximation for “genomic variance” 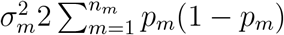 (VanRaden, 2008; Hayes *et al*., 2009) is scaled variance between marker effects and not genetic variance (Gianola *et al*., 2009; de los Campos *et al*., 2015). The scaling factor is the sum of expected variances for marker genotypes assuming Hardy-Weinberg equilibrium. Comparison of this approximation with (Eq. 1) shows that the approximation ignores linkage-disequilibrium and non-Hardy-Weinberg components of genetic variance as well as uses variance between marker effects instead of QTL effects. However, linkage-disequilibrium affects estimate of variance between marker effects. At any rate, this “misspecified” estimate of genetic variance is not useful for temporal or genomic analyses. We view variance between marker effects simply as a statistical/model parameter that facilitates model fitting to observed data. We describe the model fitting and estimation of variances in the next sub-section.

### 2.4 Statistical and computational approaches

We used the empirical and full Bayesian approach to fit the model (Eq. 2-4) with marker genotypes or their leading principal components. To fit the model (Eq. 2-4) we note that this is the ridge regression applied to marker genotype data (Whittaker *et al*., 2000; Meuwissen *et al*., 2001; de los Campos *et al*., 2013). Given the variances 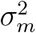 and 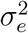 we can estimate fixed effects ***b*** and marker effects ***m*** by solving the mixed model equations:

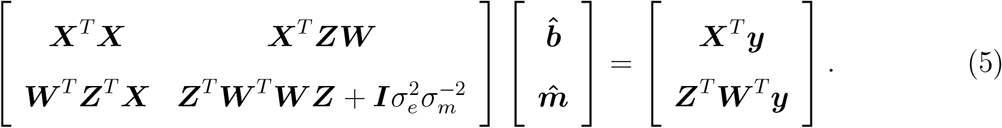

Specifically, the solution of (Eq. 5) is the conditional expectation 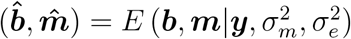. With these estimates we can obtain estimates of genetic values as 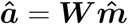. These estimates have some error and ignoring it in the framework will underestimate genetic variance. To see this, imagine we have very little phenotypic information such that marker estimates will effectively follow the prior (Eq. 3). In that case, marker estimates will effectively all equal zero and any variance calculation will return zero. As shown by Sorensen *et al*. (2001) and Lehermeier *et al*. (2017) we can account for this uncertainty by estimating genetic variances from posterior samples of genetic values or marker effects. For the model (Eq. 2-4, 5) we can obtain posterior samples from the multivariate normal distribution:

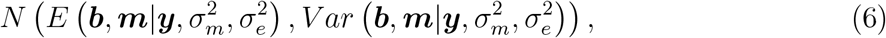

where conditional variance *Var*(***b, m***|***y***, 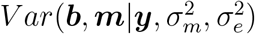) can be obtained by solving the left-hand-side of the system of equations (Eq. 5) (Sorensen and Gianola, 2007).

Once we obtained samples of marker effects from (Eq. 6) we have treated marker genotypes and marker effects respectively as QTL genotypes and QTL effects and analysed temporal and genomic trends in genetic variance as described above. Specifically, for each sample of marker effects we have estimated genetic values and their variance for each group of individuals in the breeding programme (parents, F_1_ progeny, headrows, …) across years for the temporal analysis and further partitioned across genome for the genomic analysis. This procedure gave us posterior distribution for all these variances. In results we compare how these posterior distributions match the true variances from simulation. In addition, we also calculated the continuous ranked probability score (CRPS) to compare whole posterior distributions to true values to asses both accuracy and precision and with this quantify accounting for the uncertainty of estimation. For an intuitive description of CRPS see Selle *et al*. (2019).

When variances are unknown, we can use the empirical Bayesian approach (Efron, 1996; Sorensen and Gianola, 2007) and estimate most likely variances given the data and use them to calculate conditional expectation and variance as well as draw samples from (Eq. 6). Alternatively, we can use the full Bayesian approach by specifying prior distribution for all model parameters and obtain posterior distribution 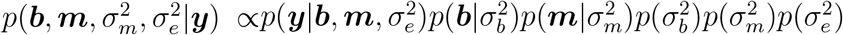 (Sorensen and Gianola, 2007).

We fitted the model (Eq. 2-4) both with the full and the empirical Bayesian approach. We first used MCMC for a full Bayesian approach and used one chain with 100,000 samples, 10,000 burn-in and saved every 100th sample to obtain 900 samples of all model parameters. For the empirical Bayesian approach, we also obtained 900 samples, but used posterior mean for the marker effect and residual variances estimated from the full Bayesian approach when sampling from (Eq. 6).

Since genomic analyses can be time-consuming we have also analysed use of approximation for marker genotypes with their leading principal components. We changed the model (Eq. 2-4) into:

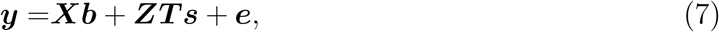

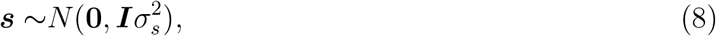

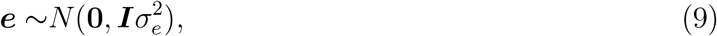

where ***T*** is an *n*_*i*_ ×*n*_*p*_ score matrix obtained from a truncated singular value decomposition of genotypes with the *n*_*p*_ leading principal components such that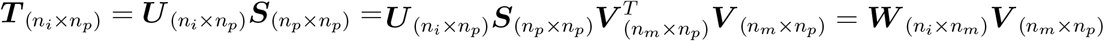, ***s*** is an *n*_*p*_ × 1 vector of *n*_*p*_ principal component effects and 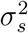 is variance between principal component effects (Hastie and Tibshirani, 2004; Tusell *et al*., 2013; Ødegård *et al*., 2018). This model is structurally the same as the model (Eq. 2-4) and we fitted it in the same way. We approximated marker effect samples by ***m***^*i*^ = ***V s***^*i*^, where ***s***^*i*^ is the *i*-th sample of principal component effects. Once we approximated marker effect samples we used the same approach as described above. We investigated different number of principal components (10, 50, 100, 500, 1000, 2000, and 3420). In our simulation these numbers of principal components respectively explained 14%, 38%, 52%, 84%, 94%, 99%, and 100% of marker genotype variation.

### 2.5 Software implementation

We have simulated the wheat breeding programme with the AlphaSimR R package (https://cran.r-project.org/web/packages/AlphaSimR/index.html) (Gaynor *et al*., 2020). We have fitted the model with the AlphaBayes software (https://www.alphagenes.roslin.ed.ac.uk/alphabayes) (Gorjanc and Hickey, 2019). We used R (R Core Team, 2019) for post-processing the AlphaBayes marker effect samples and further analyses. We used the scoringRules R package to calculate the continuous ranked probability score (CRPS) (Jordan *et al*., 2019).

## 3 Results

Overall the results show that estimates from the data following the framework were in concordance with the true values for temporal and genomic analysis. We separate the result section into three areas to facilitate presentation: (1) temporal analysis, (2) genomic analysis, and (3) computational analysis.

### 3.1 Temporal analysis

The genetic and genic variance changed through the breeding cycle. We show this in figure 3 with the true and estimated genetic and genic variances for different stages of one breeding cycle. As expected, genetic variation in F_1_ progeny across multiple crosses was lower than in the parents as this variance indicates variance in parent averages between crosses. When we generated doubled haploids for these full-sib families (HDRW stage), genetic variation was regenerated to the level in parents due to recombination and complete inbreeding. Genetic variation gradually reduced through the breeding cycle due to the selection from headrows to elite yield trial. This change was particularly evident for genetic variance, but less for genic variance. Also, genetic variance was consistently smaller than genic variance. The estimates of genetic and genic variance matched the true values well across all breeding stages. There was a larger uncertainty in the estimate of genetic variance in elite yield trial than in other stages.

**Figure 3:**
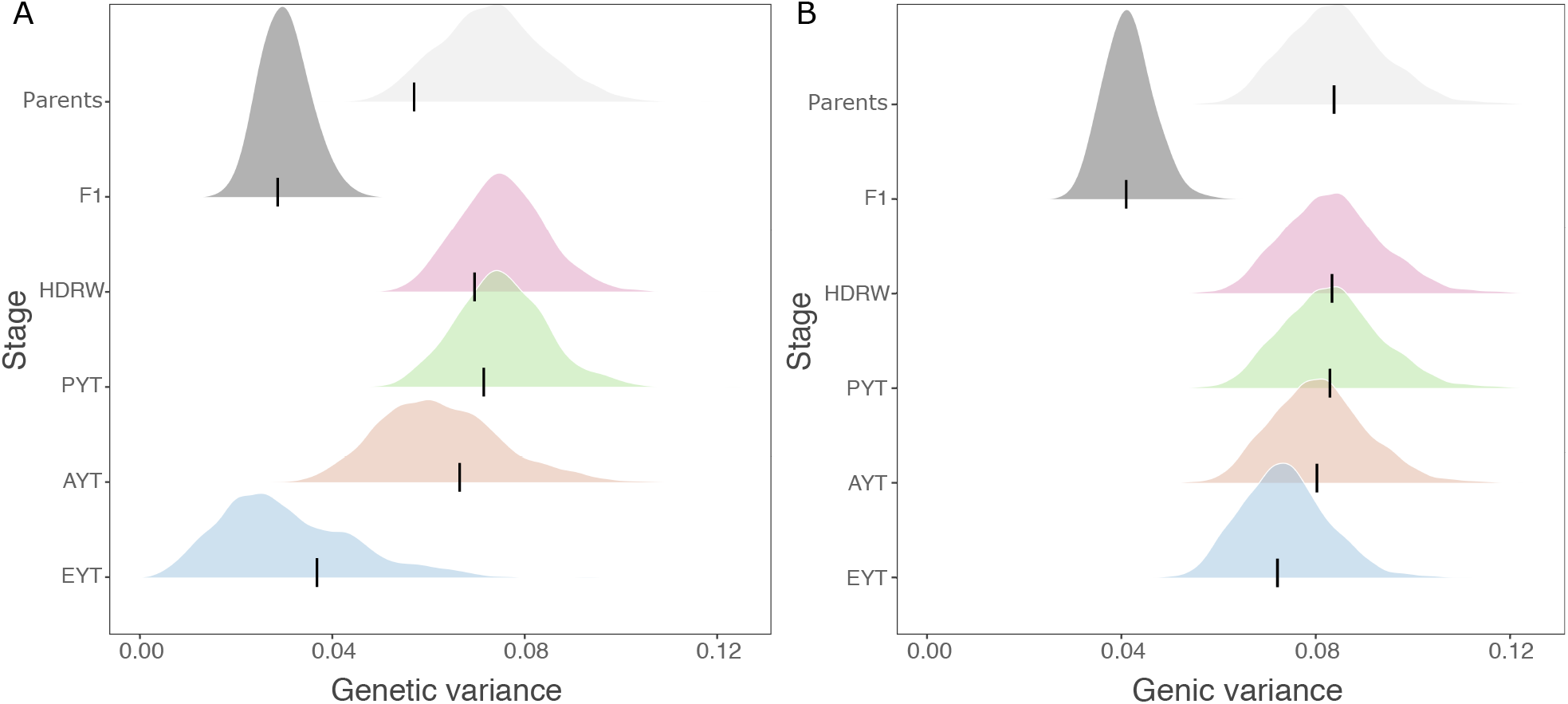
Genetic (A) and genic (B) variance estimated with the full Bayesian approach for parents in year 16, F_1_ progeny (F1) in year 17, headrows (HDRW) in year 18, preliminary yield trial (PYT) in year 19, advanced yield trial (AYT) in year 20, and elite yield trial (EYT) in year 21; black lines denote the true values and densities depict posterior distributions

Genetic variation decreased over years and genetic variance was consistently smaller as well as more variable than genic variance across years. We show this in figure 4 with the true and estimated temporal trends of genetic and genic variances for different breeding stages. Variances between the breeding stages differed as mentioned before, but in this figure we also see a consistent decrease over the years. This decrease was variable for genetic variance, but not for genic variance. This variability increased from early to late breeding stages as there was less and less individuals in a stage. The estimates of genetic and genic variance matched the true values very well across all breeding stages and years.

**Figure 4:**
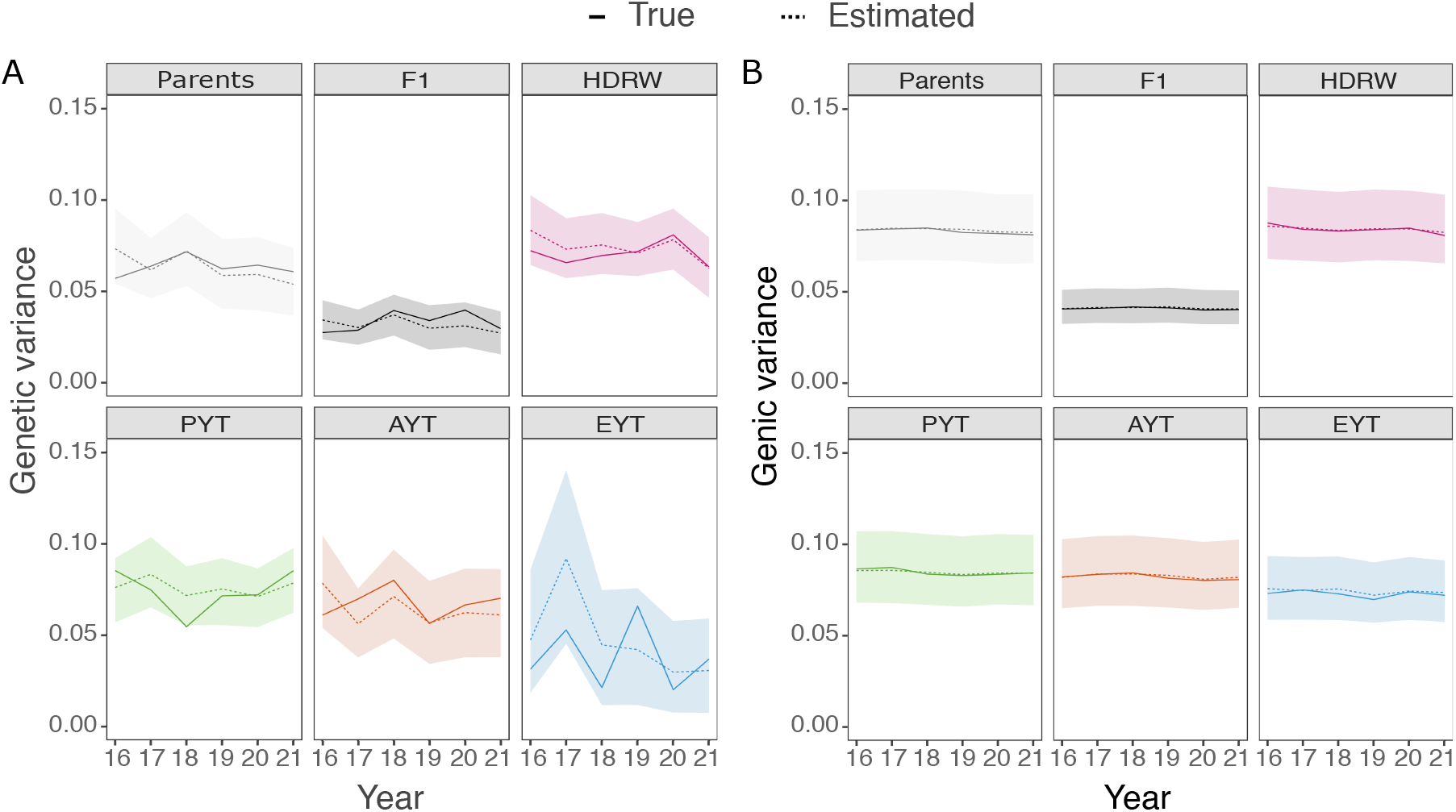
Temporal trends in genetic (A) and genic (B) variance estimated with the full Bayesian approach for parents, F_1_ progeny (F1), headrows (HDRW), preliminary yield trial (PYT), advanced yield trial (AYT), and elite yield trial (EYT); solid lines denote the true value, dashed lines denote posterior means and polygons depict 95% posterior quantiles

### 3.2 Genomic analysis

Genomic analysis enabled accurate partitioning of whole-genome genetic variance into whole-genome genic variance and whole-genome linkage-disequilibrium covariances. We show this in figure 5 with true and estimated variances and covariances for headrows and elite yield trial from one breeding cycle. The figure shows previously described differences in genetic and genic variances as well as a substantial change in the between-chromosome linkage-disequilibrium covariance, which was the main driver of change in genetic variance between headrows and the elite yield trial. Specifically, genetic variance decreased from 0.0754 in headrows in year 18 to 0.0307 in the elite yield trial in year 21, with a change of 0.0447 (59% reduction). This overall change was due to 0.01 change in genic variance (22% of the initial genetic variance), 0.0036 change in within-chromosome linkage-disequilibrium co-variance (8% of the initial genetic variance) and 0.0311 change in between-chromosome linkage-disequilibrium covariance (70% of the initial genetic variance). We again note that the estimates matched the true values well.

**Figure 5:**
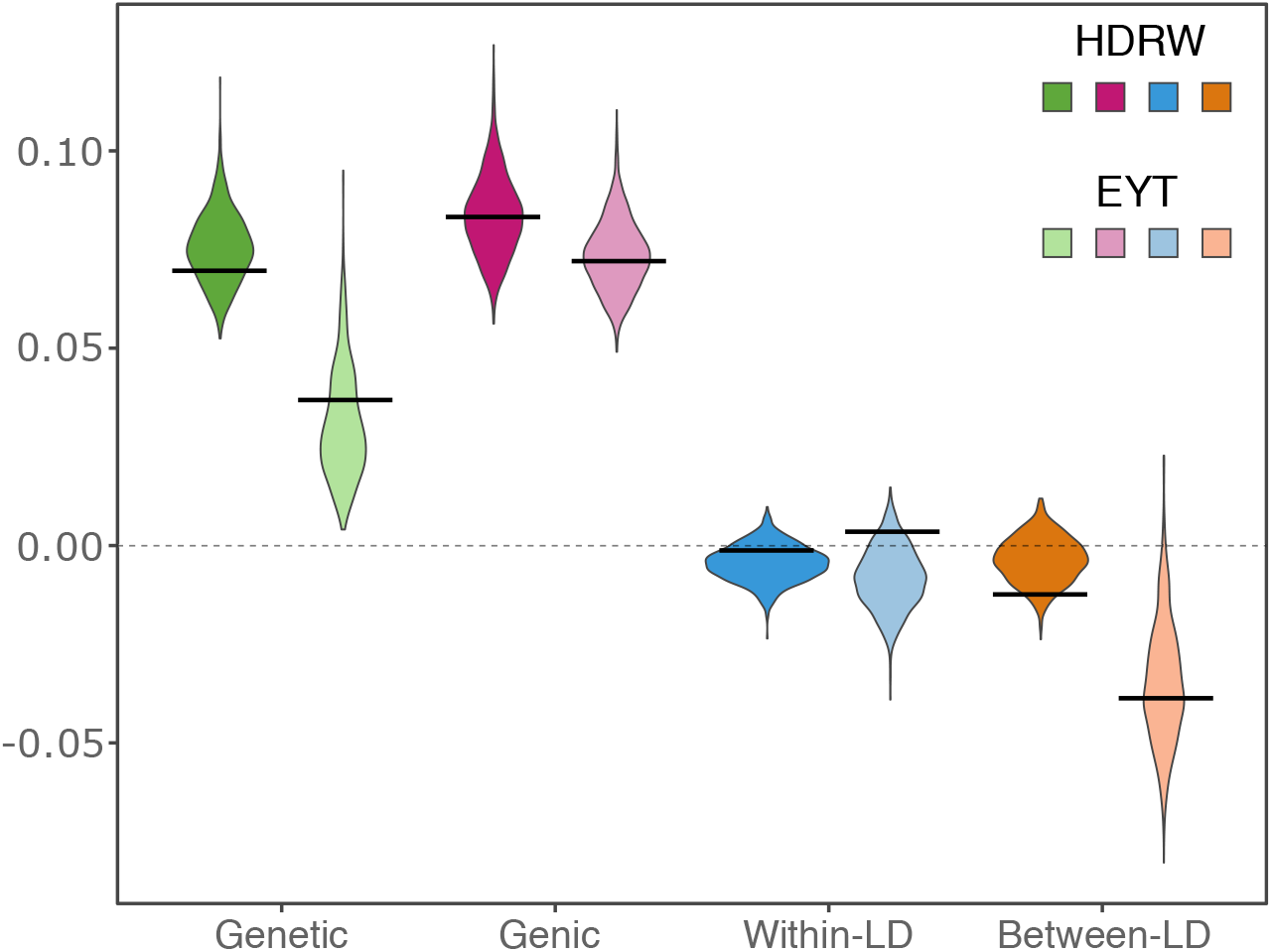
Whole-genome genetic and genic variances, and within- and between-chromosome linkage disequilibrium (LD) covariances with the full Bayesian approach for headrows (HDRW, year 18) and elite yield trial (EYT, year 21); genetic variance is the sum of genic variance, within- and between-chromosome LD (see Fig. 2); black lines denote true values and violins depict posterior distributions

Genomic analysis enabled also accurate partitioning of whole-genome genetic variance for specific chromosomes. We show this in the supplementary material with a series of tables (S1-S4) and one figure (S1). The tables show genetic variance and its components (genic variance, within-chromosome linkage-disequilibrium covariance and between-chromosome linkage-disequilibrium covariance) by 21 chromosomes as well as how these values add up to the whole-genome variance. We show this partitioning for QTL genotypes (Table S1), marker genotypes (Table S2), true genetic values (Table S3), and estimated genetic values (Table S4). The figure S1 compares the true and estimated genetic values directly. The aim of this supplementary material is to demonstrate how we estimate variation in true genetic values, which is driven by unknown QTL and unknown QTL effects, by using marker genotypes and estimated marker effects. We make five observations. First, the analysis of QTL genotypes showed that whole-genome and chromosome genetic variance in unselected headrows is largely driven by genic variance, but there are some chromosomes with a substantial within-chromosome or between-chromosome linkage-disequilibrium covariance. Second, the magnitude of linkage-disequilibrium covariances increased in the elite yield trial, which reduced the whole-genome genetic variance. However, between-chromosome linkage-disequilibrium was larger than within-chromosome linkage-disequilibrium. Third, the analysis of marker genotypes followed the same trends, but the values were sustainability larger due to larger number of markers than QTL. Fourth, the analysis of true genetic values resulted in much smaller values for variances than the analysis of QTL genotypes because most QTL have small effects, but the relative magnitude of variation and its partitioning was similar. Fifth, the analysis of estimated genetic values followed closely the analysis of true genetic values - most deviations were observed for the elite yield trial, but all posterior distributions encompassed the true value. This analysis pertains to one single dataset to show that estimates are reasonable for a specific dataset.

### 3.3 Computational analysis

Full and empirical Bayesian approaches had similar posterior mean estimates of variances, but empirical Bayesian approach had smaller posterior standard deviation. We show this in figure 6 with a comparison of posterior means and posterior standard deviations for genetic and genic variance between the two approaches. The posterior means matched well for both types of variances. The posterior standard deviation was smaller with the empirical Bayesian approach, in particular for the genic variance. Comparison with the true values however showed good concordance with the empirical Bayesian posterior means (Fig. S2 and S3).

**Figure 6:**
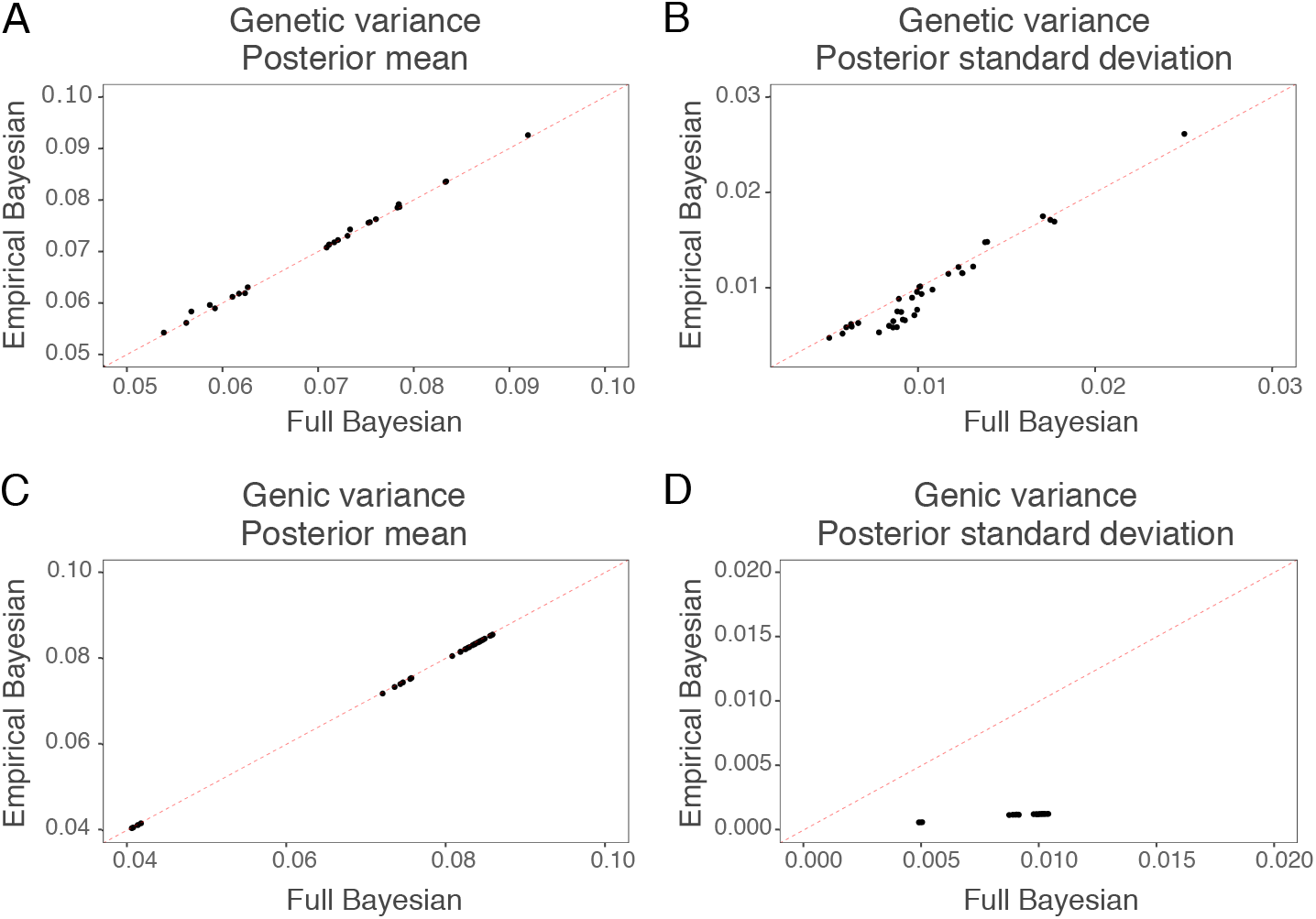
The empirical Bayesian approach versus the full Bayesian approach for posterior mean of genetic variance (A), posterior mean of genic variance (B), posterior standard deviation of genetic variance (C), and posterior standard deviation of genic variance (D); equal value is represented by the dashed red line

Additional evaluation with multiple replicates showed that the full and empirical Bayesian results were consistently estimated for genetic and genic variance estimates. We show this in table 1 with continuous ranked probability score (CRPS) of genetic and genic variances for full and empirical Bayesian approaches by breeding stage. Note that CRPS is negatively oriented - lower values indicate better estimate compared to the true value in terms of accuracy and precision. CRPS for genetic variance matched closely between the full and empirical Bayesian approaches. On the other hand, they differ more for genic variance, with better (lower) values for the full Bayesian approach, albeit there was large variability across years and replicates. CRPS was larger (worse) for genic variance than for genetic variance.

**Table 1:** Continuous ranked probability score (CRPS × 1000 - lower is better: mean ± standard deviation over six years and ten replicates) for genetic and genic variance estimated by the full Bayesian and the empirical Bayesian for parents, F_1_ progeny, headrows (HDRW), preliminary yield trial (PYT), advanced yield trial (AYT), and elite yield trial (EYT)

| Stage | Genetic |  | Genic |  |
| --- | --- | --- | --- | --- |
|  | Full | Empirical | Full | Empirical |
| Parents | $59 \pm 40$ | $60 \pm 41$ | $300 \pm 93$ | $351 \pm 97$ |
| $F_1$ | $42 \pm 39$ | $42 \pm 40$ | $40 \pm 44$ | $48 \pm 52$ |
| HDRW | $45 \pm 32$ | $46 \pm 37$ | $297 \pm 94$ | $348 \pm 99$ |
| PYT | $63 \pm 57$ | $64 \pm 64$ | $296 \pm 94$ | $348 \pm 98$ |
| AYT | $66 \pm 63$ | $66 \pm 64$ | $294 \pm 92$ | $344 \pm 97$ |
| EYT | $79 \pm 45$ | $80 \pm 46$ | $70 \pm 75$ | $84 \pm 90$ |

Approximation with leading principal components accurately estimated genetic variance when we used sufficient number of principal components, but this was never the case for genic variance. We show this in figure 7 with estimation error, defined as the difference between the true and estimated value, for genetic and genic variance as a function of the number of leading principal components. The estimation error decreased as we increased the number of leading principal components. It decreased quickly for the genetic variance - there was no error once we captured about 80% of variation in marker genotypes. In our simulated dataset we achieved this with 500 leading principal components. On the other hand, the estimation error decreased slowly for the genic variance and we never recovered the true estimate, even if we used all the principal components. The estimation error was smallest in the F_1_ pogeny, followed by the elite yield trial, while the largest estimation error were in headrows.

**Figure 7:**
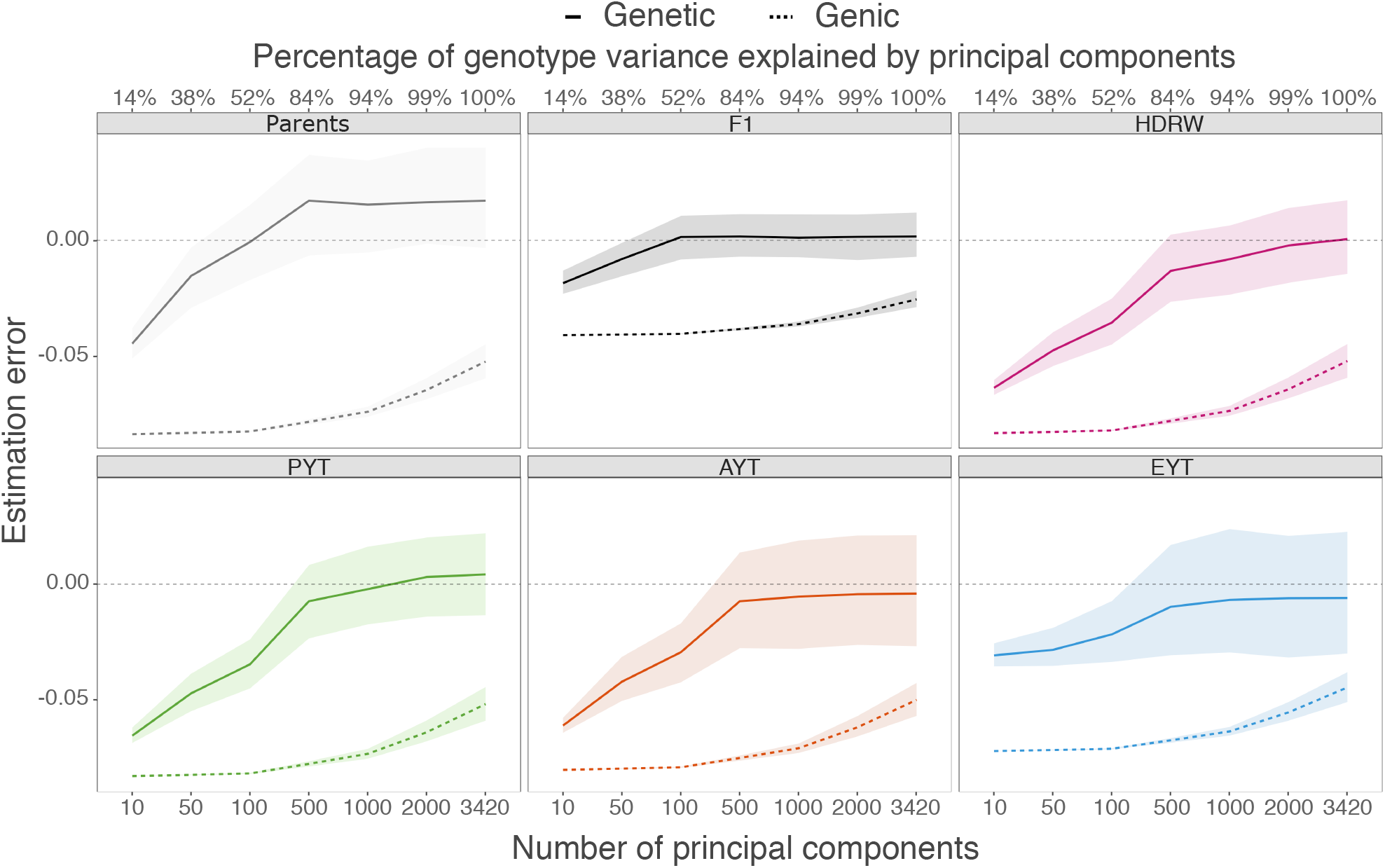
Estimation error in genetic and genic variances as a function of the number of principal components in parents in year 16, F_1_ progeny (F1) in year 17, headrows (HDRW) in year 18, preliminary yield trial (PYT) in year 19, advanced yield trial (AYT) in year 20, and elite yield trial (EYT) in year 21; horizontal dashed line represents no estimation error

## 4 Discussion

The results show that the framework for temporal and genomic analysis of genetic variation is flexible, accurate and enables assessing the sustainability of a breeding programme as well as processes that change genetic variance. These results highlights four topics for discussion in line with the structure of results: (1) temporal analysis of genetic variance, (2) genomic analysis of genetic variance, (3) computational aspects and (4) assumptions of this study.

### 4.1 Temporal analysis

This study will help breeders to assess the amount of genetic variance in their programmes and with this better management of its utilization for future genetic gains. Genetic variance (specifically its square root) is key component of the breeders equation for predicting response to selection (Lush, 1937; Falconer and Mackay, 1996). While breeding programmes routinely estimate genetic variance for traits under selection, most estimates pertain to a group of individuals that is arguably not the most relevant for routine breeding (Piepho *et al*., 2008). Specifically, with the pedigree-based model the estimate of genetic variance pertains to pedigree founders, which can be several generations removed from currently interesting individuals. Further, pedigree founders often span multiple generations due to incomplete pedigrees and as such the corresponding estimate of genetic variance does not have a clearly defined time point. Estimates of genetic variance from genome-based models pertains to all genotyped individuals, which again does not have a clearly defined time point. In addition, the “genomic variance” is plagued with model “misspecification” (Gianola *et al*., 2009; de los Campos *et al*., 2015), see also Schreck *et al*. (2019).

The proposed framework that builds on the work of Sorensen *et al*. (2001), Lehermeier *et al*. (2017) and Allier *et al*. (2019) enables straightforward temporal analysis both in terms of years and stages of a breeding programme. The framework uses all the available data spanning multiple years (generations) to estimate model parameters, which are in turn used to infer genetic values and their variances. Such flexibility of using all data but producing estimates for any group of individuals is crucial to inform breeders how much genetic variance they have at hand so that they can react accordingly. For example, temporal trends in genetic and genic variance enable straightforward trait specific estimation of effective population size (Gorjanc *et al*., 2018). Using this approach in this study we estimated effective population size for the parents at 111. This estimate suggests that the simulated breeding programme is sustainable (Falconer and Mackay, 1996; Hill, 2016; Lynch and Walsh, 1998; Walsh and Lynch, 2018) as indicated by small changes in genetic variance between years. Possible reactions to a temporal analysis by a breeder could be keeping the current breeding programme as it is, implementing active management of genetic variance using techniques such as optimal contribution selection (e.g., Woolliams *et al*., 2015; Akdemir and Sánchez, 2016; Gorjanc *et al*., 2018; Akdemir *et al*., 2019), germplasm exchange with other programmes or in the extreme introgressing landrace germplasm (e.g., Gorjanc *et al*., 2016).

There are also other approaches to temporal analysis of genetic variance. Tsuruta *et al*. (2004) used the random regression model to model genetic values and their variance over years and Hidalgo *et al*. (2020) used sliding time intervals in the same fashion. Both methods have some drawbacks - random regression can be computationally expensive, while time intervals must be sufficiently large to obtain accurate estimates. These two approaches respectively enrich the model or slice the data to estimate genetic variances over time, while the proposed framework treats model variance parameters and genetic variances over time as two separate sets. We will address these differences at the end of discussion. Hidalgo *et al*. (2020) used sliding time intervals to investigate changes in genetic (co)variances for a breeding programme that recently implemented genomic selection. They observed rapid changes in genetic (co)variances with the implementation of genomic selection. Their results clearly highlight a need for breeder’s reaction and further investigation. One such investigation should be on which components of genetic variance changed with the implementation of genomic selection.

### 4.2 Genomic analysis

The proposed framework can estimate size and trends for genomic components of genetic variance. We have followed a standard quantitative genetics decomposition of genetic variance (Bulmer, 1971; Lynch and Walsh, 1998; Gianola *et al*., 2009; Walsh and Lynch, 2018), which involves a component due to variance of genotypes and their allele substitution effects at every quantitative trait locus (genic variance) and a component due to covariance between genotypes and their allele substitution effects between loci on one chromosome (within-chromosome linkage-disequilibrium covariance) and between chromosomes (between-chromosome linkage-disequilibrium covariance). Our results show promising utility of the proposed framework. We showed this decomposition for quantitative trait locus genotypes, marker genotypes, true genetic values and estimated values, all at the whole-genome and chromosome level. These results confirmed the prediction of Bulmer (1971) that directional selection on total genetic values or their functions (phenotype) induces negative linkage-disequilibrium and that this component can cause rapid and major changes in genetic variance (Lynch and Walsh, 1998; Walsh and Lynch, 2018). We note that this negative linkage-disequilibrium is a function of genotype combinations between loci as well as their allele substitution effects. Therefore, we have to distinguish between linkage-disequilibrium between genotypes, which is trait agnostic, and linkage-disequilibrium between locus genetic values (see Tables S1-S4).

The importance of linkage-disequilibrium in estimating genetic variance with genomic data is growing (de los Campos *et al*., 2015; Lehermeier *et al*., 2017; Allier *et al*., 2019). Our study added to this literature with a simulation study and demonstrating temporal changes in linkage-disequilibrium under selection both within one breeding cycle (headrows to elite yield trial) and between breeding cycles over years. We observed larger changes within breeding cycles than between, which can be explained by strong selection within cycles and recombinations among initial parent genomes between cycles. Interestingly, we observed large between-chromosome linkage-disequilibrium covariance in comparison to within-chromosome. This is at odds with physical linkage between loci within a chromosome and no such linkage between loci on separate chromosomes. Our explanation for this is that there is a larger number of combinations between loci on separate chromosomes than within chromosomes. Further, limited recombination constrains selection to induce linkage-disequilibrium within chromosomes compared to between chromosomes. To put this into perspective, in the analysed example we observed a 59% change in genetic variance within a breeding cycle (headrows to elite yield trial) of which 22% was due to the change in genic variance, 8% was due to the change in within-chromosome linkage-disequilibrium covariance and 70% was due to the change in between-chromosome linkage-disequilibrium covariance. These overall values varied considerably between chromosomes, where we emphasise that our simulation randomly placed loci and randomly allocated effects from one common distribution. These assumptions are likely too simple and indeed Allier *et al*. (2019) observed strong variation between chromosomes in maize. All in all, these results indicate that linkage-disequilibrium is an important component of the genetic variance in line with the theoretical work of Bulmer (1971) and Mather and Jinks (2013).

We expected that we will underestimate genic variance in this breeding study, but have not observed this. We have simulated breeding programme with directional selection, which induces negative linkage-disequilibrium (Bulmer, 1971) due to repulsion linkage (Mather and Jinks, 2013). We expected that repulsion linkage will “hide” variation in some genome regions due to a lack of variation in haplotypes and that we will therefore underestimate genic variance. This did not happen either because effective population was reasonably large (111), selection was not too strong or there were sufficient number of markers. However, across multiple replicates the continuous ranked probability score was worse for genic than genetic variance, which could indicate this systematic underestimation.

The presented framework for genomic analysis of genetic variance will pave the way for analysing processes that change the variance. While selection induces linkage-disequilibrium between loci it also changes allele frequencies (Bulmer, 1971; Lynch and Walsh, 1998; Gorjanc *et al*., 2015; Walsh and Lynch, 2018). Another important process is drift, which is always present in breeding programmes due to small effective population sizes. Distinguishing between selection and drift in such populations is difficult (Lynch and Walsh, 1998; Gorjanc *et al*., 2015; Walsh and Lynch, 2018) and further work is required. Similarly, population structure and admixture between populations can influence genetic variance and should be addressed in the future. One way to treat population structure would be to partition individuals by sub-population and calculate separate genetic variances as well as covariances between sub-populations. This approach breaks down with admixture. Admixture could be approached by using whole population genome trees with recombination (Kelleher *et al*., 2019) and label individuals and genome segments with originating sub-populations and expand the framework into population analysis of genetic variance.

A final note on genomic analysis is that the proposed framework does not depend on the assumption of Hardy-Weinberg and linkage equilibrium. It is common to see expressions for genetic variance at a locus of the form 2*p*(1 − *p*)*α*^2^, which assumes independent binomial sampling of alleles with probability *p* (Hardy-Weinberg equilibrium). In some breeding programmes there is an excess of homozygotes over heterozygotes, particularly in plant breeding programmes that use selfing. In this case we have a clear deviation from the Hardy-Weinberg equilibrium and the expression 2*p*(1 − *p*)*α*^2^ will underestimate genetic variance. To see this consider *p* = 0.5 and *α* = 1, which gives 2*p*(1 − *p*)*α*^2^ = 0.5, but if we only have reference and alternative homozygotes (50% each) the actual variance is doubled due to complete inbreeding (Wright, 1931). While there are expressions that involve inbreeding 2*p*(1 − *p*)(1 + *F*)*α*^2^, where 2*p*(1 − *p*)(1 + *F*) is variance of genotypes under non-random mating, we suggest a simpler straightforward calculation of sample variance of genotypes at a locus and multiplying that variance with *α*^2^. Bulmer (1976) was aware of these differences and partitioned genic variance into the value expected under Hardy-Weinberg equilibrium (binomial sampling of alleles) 2*p*(1 − *p*)*α*^2^ and deviation due to non-random mating *Fα*^2^.

### 4.3 Computational aspects

The proposed framework is based on Sorensen *et al*. (2001), Lehermeier *et al*. (2017), and Allier *et al*. (2019) that used the full Bayesian approach and MCMC sampling. We performed our analyses with the full and empirical Bayesian approach and found a good concordance between the two approaches and true values. However, there was tendency of the empirical Bayesian approach to underestimate uncertainty of inferred genetic variances, due to ignoring uncertainty in estimating model variance parameters. This is expected, but it seems that the difference is not large, though this will vary between datasets. The full Bayesian analysis with marker-based models is not too computationally demanding if the number of markers is not too large (10-50K markers can be handled with ease). The full Bayesian analysis can be quite demanding with genome-based model on individuals if the number of individuals is large, but equivalence with the marker-based model means we can fit one or another model and back-solve desired effects (Strandén and Garrick, 2009). There are also frequentist approaches that account for uncertainty of estimating variance components (e.g. Kenward and Roger, 1997). For the genomic analysis there is an advantage (in terms of flexibility) in working with marker effects and marker genotypes.

The observation that leading principal components underestimate genic variance require further studies. We expected that increasing the number of leading principal components will reduce the estimation error, which we observed for genetic variance, while we observed consistent underestimation for genic variance - even with all principal components. Since we had more markers than individuals this is likely due to the fact that “null” components would still have some uncertainty in estimation, which we ignored and therefore underestimated genic variance. Methods presented in the supplementary of Listgarten *et al*. (2012) could be used to correct for this.

### 4.4 Assumptions

In this study we made two related assumptions and one unrelated assumption. First, we assumed that allele effects are constant over time and across groups of individuals. This is a reasonable assumption in a sense that we used all the available data to accurately estimate marker effects. Time- or background-specific estimation could better reflect reality, because linkage-disequilibrium is changing over time, but getting accurate estimates from less data is challenging and so is defining time intervals or backgrounds. The random regression and time interval approaches (Tsuruta *et al*., 2004; Hidalgo *et al*., 2020) have an advantage with this aspect, but a limitation in terms of flexibility for the genomic analysis of genetic variance. This aspect of variable effects will likely be more important with breeding programmes that introgress germplasm from other populations, but there will also likely be too little data to estimate separate effects. Estimation of background-specific effects is an active research area in genetics with growing datasets across various populations (e.g., Peterson *et al*., 2019; van den Berg *et al*., 2020). Second, we assumed fully additive genetic architecture under which allele effects are constant across time and groups of individuals. While both theory and data indicate that average effect of an allele substitution capture majority of genetic variance (Hill *et al*., 2008), recognition of dominance and epistasis is growing (e.g., Varona *et al*., 2018). Recognition of genotype interactions with environment is also growing (e.g., Tolhurst *et al*., 2019). The proposed framework can be expanded to these settings, but the success of inferring various variances, potentially in different environments, will critically depend on volume of data to estimate much larger number of parameters. Third, we assumed a sufficiently dense panel of markers that collectively closely track quantitative trait loci. Insufficient number of markers will deteriorate the ability of the proposed framework to capture genetic variance at and between quantitative trait loci.

## Acknowledgments

The authors acknowledge the financial support from the BBSRC ISP to The Roslin Institute BBS/E/D/30002275, the grant BBSRC IIA PIII-036 and the University of Edinburghs Data-Driven Innovation Chancellors fellowship. This work has made use of the resources provided by the University of Edinburgh Compute and Data Facility (ECDF) (http://www.ecdf.ed.ac.uk).

## Data Availability

We provide all analysis scripts at: https://git.ecdf.ed.ac.uk/HighlanderLab_public/llara_gen_var_plants.

## Supplementary Material

**Table S1:** Genetic variance partitioned into genic variance and within- and between-chromosome linkage-disequilibrium (LD) covariances by chromosome for **QTL genotypes** in headrows (HDRW, year 18) and elite yield trial (EYT, year 21); the genetic variance is the sum of genic variance, within-LD and between-LD (see Fig. 2)

| Chr | HDRW |  |  |  | EYT |  |  |  |
| --- | --- | --- | --- | --- | --- | --- | --- | --- |
|  | Genetic | Genic | Within-LD | Between-LD | Genetic | Genic | Within-LD | Between-LD |
| 1 | 98.2 | 61.9 | 36.3 | -5.1 | 91.6 | 55.4 | 36.2 | 40.6 |
| 2 | 49.9 | 55.3 | -5.4 | -7.4 | 65.0 | 44.5 | 20.5 | -39.2 |
| 3 | 50.5 | 55.8 | -5.3 | -15.5 | 52.6 | 47.4 | 5.2 | -97.4 |
| 4 | 48.0 | 52.5 | -4.5 | -7.9 | 20.6 | 51.4 | -30.8 | 14.2 |
| 5 | 49.8 | 55.6 | -5.8 | -17.4 | 41.6 | 43.0 | -1.4 | -51.6 |
| 6 | 55.1 | 60.4 | -5.3 | 1.8 | 45.2 | 57.2 | -12.0 | -58.6 |
| 7 | 53.2 | 61.9 | -8.6 | 0.1 | 32.2 | 54.0 | -21.8 | 40.4 |
| 8 | 84.6 | 47.7 | 36.9 | 7.5 | 153.0 | 43.0 | 110.0 | 153.2 |
| 9 | 73.8 | 65.7 | 8.2 | -52.4 | 47.8 | 58.3 | -10.5 | -38.6 |
| 10 | 65.1 | 57.5 | 7.6 | 8.0 | 104.6 | 57.2 | 47.4 | -281.4 |
| 11 | 49.6 | 61.5 | -11.9 | -1.0 | 25.6 | 58.8 | -33.1 | -14.6 |
| 12 | 40.9 | 62.4 | -21.5 | 4.4 | 89.8 | 51.2 | 38.6 | -82.4 |
| 13 | 48.2 | 63.4 | -15.2 | 14.0 | 39.4 | 49.2 | -9.8 | 9.8 |
| 14 | 68.4 | 59.2 | 9.3 | 14.6 | 34.6 | 48.4 | -13.8 | -1.6 |
| 15 | 50.2 | 56.8 | -6.6 | -2.2 | 55.2 | 48.2 | 7.0 | 69.6 |
| 16 | 86.6 | 61.6 | 24.9 | -48.8 | 73.2 | 50.6 | 22.6 | -35.0 |
| 17 | 65.0 | 58.5 | 6.6 | 29.5 | 74.0 | 55.8 | 18.2 | 134.6 |
| 18 | 57.0 | 60.0 | -3.0 | -10.5 | 53.0 | 49.9 | 3.0 | -24.2 |
| 19 | 54.9 | 60.7 | -5.8 | 4.7 | 37.4 | 50.0 | -12.6 | 7.0 |
| 20 | 29.6 | 58.2 | -28.6 | 8.9 | 36.4 | 47.0 | -10.6 | 19.0 |
| 21 | 34.3 | 58.7 | -24.4 | -1.5 | 28.0 | 53.9 | -25.9 | -13.2 |
| Sum | 1213.1 | 1235.1<sup>1</sup> | -22.0<sup>2</sup> | -76.2<sup>3</sup> | 1200.8 | 1074.4<sup>1</sup> | 126.3<sup>2</sup> | -249.4<sup>3</sup> |
| Whole-genome<sup>1+2+3</sup> | 1136.9 |  |  |  | 951.4 |  |  |  |

**Table S2:** Genetic variance partitioned into genic variance and within- and between-chromosome linkage-disequilibrium (LD) covariances by chromosome for **marker genotypes** in headrows (HDRW, year 18) and elite yield trial (EYT, year 21); the genetic variance is the sum of genic variance, within-LD and between-LD (see Fig. 2)

| Chr | HDRW |  |  |  | EYT |  |  |  |
| --- | --- | --- | --- | --- | --- | --- | --- | --- |
|  | Genetic | Genic | Within-LD | Between-LD | Genetic | Genic | Within-LD | Between-LD |
| 1 | 286.9 | 310.2 | -23.3 | -156.1 | 151.2 | 278.7 | -127.4 | 619.1 |
| 2 | 383.6 | 288.6 | 95.0 | -151.0 | 450.4 | 246.4 | 204.0 | 18.8 |
| 3 | 270.1 | 289.9 | -19.8 | 44.2 | 435.2 | 257.4 | 177.8 | 829.2 |
| 4 | 371.8 | 288.4 | 83.4 | 125.8 | 268.0 | 267.0 | 1.0 | 507.2 |
| 5 | 317.5 | 286.2 | 31.3 | 20.6 | 117.4 | 211.1 | -93.7 | -24.0 |
| 6 | 347.0 | 290.8 | 56.2 | 59.1 | 395.4 | 278.9 | 116.5 | 848.3 |
| 7 | 337.1 | 311.9 | 25.2 | -172.7 | 692.8 | 289.4 | 403.4 | -1021.2 |
| 8 | 340.5 | 274.2 | 66.4 | -243.9 | 263.8 | 221.6 | 42.2 | -1231.2 |
| 9 | 290.1 | 302.8 | -12.6 | 11.7 | 133.6 | 285.6 | -151.9 | 242.4 |
| 10 | 403.9 | 317.0 | 86.9 | -16.6 | 473.0 | 305.2 | 167.8 | 816.6 |
| 11 | 192.7 | 304.2 | -111.4 | 45.9 | 48.6 | 290.2 | -241.6 | -129.7 |
| 12 | 316.0 | 300.9 | 15.1 | -43.6 | 230.6 | 243.5 | -13.0 | -180.2 |
| 13 | 303.6 | 294.8 | 8.8 | -175.6 | 114.6 | 245.9 | -131.4 | 416.7 |
| 14 | 285.6 | 315.7 | -30.1 | 34.5 | 95.6 | 277.6 | -182.0 | -346.5 |
| 15 | 221.1 | 292.8 | -71.8 | -32.2 | 319.2 | 256.8 | 62.5 | 25.1 |
| 16 | 396.9 | 298.3 | 98.6 | -0.2 | 215.4 | 248.4 | -33.0 | 213.4 |
| 17 | 322.9 | 301.3 | 21.7 | -24.8 | 467.4 | 283.0 | 184.3 | -1384.9 |
| 18 | 229.8 | 290.1 | -60.3 | -32.3 | 105.2 | 245.6 | -140.5 | 532.7 |
| 19 | 225.4 | 307.3 | -81.9 | 48.6 | 88.2 | 273.5 | -185.4 | -16.1 |
| 20 | 404.2 | 296.3 | 107.9 | -58.4 | 234.4 | 245.7 | -11.3 | 175.2 |
| 21 | 205.9 | 286.7 | -80.8 | -119.4 | 146.8 | 255.4 | -108.6 | 3.1 |
| Sum | 6452.6 | 6248.3<sup>1</sup> | 204.3<sup>2</sup> | -836.7<sup>3</sup> | 5446.8 | 5507.0<sup>1</sup> | -60.2<sup>2</sup> | 914.1<sup>3</sup> |
| Whole-genome<sup>1+2+3</sup> | 5615.9 |  |  |  | 6360.8 |  |  |  |

**Table S3:** Genetic variance partitioned into genic variance and within- and between-chromosome linkage-disequilibrium (LD) covariances by chromosome for **true genetic values** in headrows (HDRW, year 18) and elite yield trial (EYT, year 21); the genetic variance is the sum of genic variance, within-LD and between-LD (see Fig. 2)

| Chr | HDRW |  |  |  | EYT |  |  |  |
| --- | --- | --- | --- | --- | --- | --- | --- | --- |
|  | Genetic | Genic | Within-LD | Between-LD | Genetic | Genic | Within-LD | Between-LD |
| 1 | 0.0036 | 0.0039 | -0.0003 | -0.0010 | 0.0014 | 0.0031 | -0.0017 | -0.0056 |
| 2 | 0.0047 | 0.0046 | 0.0001 | -0.0020 | 0.0030 | 0.0033 | -0.0003 | -0.0021 |
| 3 | 0.0035 | 0.0042 | -0.0007 | 0.0011 | 0.0014 | 0.0040 | -0.0027 | 0.0028 |
| 4 | 0.0029 | 0.0039 | -0.0010 | -0.0002 | 0.0030 | 0.0036 | -0.0005 | 0.0017 |
| 5 | 0.0050 | 0.0037 | 0.0013 | -0.0008 | 0.0040 | 0.0027 | 0.0013 | 0.0004 |
| 6 | 0.0030 | 0.0026 | 0.0004 | -0.0017 | 0.0016 | 0.0025 | -0.0009 | 0.0004 |
| 7 | 0.0041 | 0.0041 | 0.0000 | 0.0002 | 0.0042 | 0.0035 | 0.0008 | -0.0002 |
| 8 | 0.0023 | 0.0035 | -0.0012 | -0.0006 | 0.0031 | 0.0036 | -0.0005 | 0.0021 |
| 9 | 0.0044 | 0.0043 | 0.0001 | -0.0002 | 0.0042 | 0.0038 | 0.0004 | -0.0040 |
| 10 | 0.0025 | 0.0033 | -0.0008 | -0.0003 | 0.0045 | 0.0033 | 0.0013 | -0.0075 |
| 11 | 0.0023 | 0.0037 | -0.0014 | 0.0004 | 0.0016 | 0.0035 | -0.0019 | -0.0052 |
| 12 | 0.0054 | 0.0043 | 0.0010 | 0.0000 | 0.0048 | 0.0036 | 0.0012 | -0.0031 |
| 13 | 0.0056 | 0.0037 | 0.0019 | -0.0005 | 0.0076 | 0.0028 | 0.0048 | -0.0087 |
| 14 | 0.0026 | 0.0045 | -0.0019 | -0.0004 | 0.0037 | 0.0039 | -0.0002 | -0.0084 |
| 15 | 0.0044 | 0.0034 | 0.0010 | -0.0004 | 0.0035 | 0.0034 | 0.0001 | 0.0001 |
| 16 | 0.0058 | 0.0053 | 0.0005 | -0.0027 | 0.0082 | 0.0042 | 0.0040 | -0.0075 |
| 17 | 0.0060 | 0.0051 | 0.0009 | -0.0019 | 0.0075 | 0.0052 | 0.0022 | 0.0008 |
| 18 | 0.0038 | 0.0042 | -0.0004 | 0.0010 | 0.0034 | 0.0032 | 0.0002 | 0.0003 |
| 19 | 0.0039 | 0.0038 | 0.0001 | -0.0020 | 0.0022 | 0.0030 | -0.0007 | 0.0038 |
| 20 | 0.0033 | 0.0036 | -0.0003 | 0.0002 | 0.0009 | 0.0026 | -0.0017 | -0.0007 |
| 21 | 0.0030 | 0.0037 | -0.0007 | -0.0006 | 0.0017 | 0.0033 | -0.0016 | 0.0019 |
| Sum | 0.0820 | 0.0833<sup>1</sup> | -0.0013<sup>2</sup> | -0.0124<sup>3</sup> | 0.0756 | 0.0721<sup>1</sup> | 0.0035<sup>2</sup> | -0.0387<sup>3</sup> |
| Whole-genome<sup>1+2+3</sup> | 0.0696 |  |  |  | 0.0369 |  |  |  |

**Table S4:** Genetic variance partitioned into genic variance and within- and between-chromosome linkage-disequilibrium (LD) covariances by chromosome for **estimated genetic values** (with the full Bayesian approach) in headrows (HDRW, year 18) and elite yield trial (EYT, year 21); the genetic variance is the sum of genic variance, within-LD and between-LD (see Fig. 2)

| Chr | HDRW |  |  |  | EYT |  |  |  |
| --- | --- | --- | --- | --- | --- | --- | --- | --- |
|  | Genetic | Genic | Within-LD | Between-LD | Genetic | Genic | Within-LD | Between-LD |
| 1 | 0.0037 | 0.0041 | -0.0004 | 0.0003 | 0.0041 | 0.0037 | 0.0004 | -0.0029 |
| 2 | 0.0034 | 0.0038 | -0.0004 | 0.0005 | 0.0031 | 0.0033 | -0.0002 | -0.0009 |
| 3 | 0.0044 | 0.0039 | 0.0005 | 0.0012 | 0.0040 | 0.0035 | 0.0005 | -0.0006 |
| 4 | 0.0033 | 0.0038 | -0.0005 | -0.0007 | 0.0035 | 0.0035 | -0.0001 | -0.0029 |
| 5 | 0.0044 | 0.0039 | 0.0005 | -0.0004 | 0.0030 | 0.0028 | 0.0001 | -0.0024 |
| 6 | 0.0037 | 0.0039 | -0.0002 | -0.0011 | 0.0027 | 0.0037 | -0.0010 | -0.0009 |
| 7 | 0.0037 | 0.0042 | -0.0005 | -0.0005 | 0.0027 | 0.0039 | -0.0011 | -0.0016 |
| 8 | 0.0031 | 0.0037 | -0.0006 | -0.0004 | 0.0023 | 0.0030 | -0.0007 | -0.0011 |
| 9 | 0.0039 | 0.0040 | -0.0001 | 0.0004 | 0.0038 | 0.0038 | 0.0000 | -0.0021 |
| 10 | 0.0037 | 0.0042 | -0.0005 | 0.0000 | 0.0030 | 0.0041 | -0.0011 | -0.0018 |
| 11 | 0.0037 | 0.0040 | -0.0003 | 0.0001 | 0.0040 | 0.0039 | 0.0002 | -0.0031 |
| 12 | 0.0041 | 0.0041 | 0.0000 | -0.0005 | 0.0038 | 0.0033 | 0.0004 | -0.0025 |
| 13 | 0.0045 | 0.0040 | 0.0005 | 0.0008 | 0.0028 | 0.0033 | -0.0005 | -0.0011 |
| 14 | 0.0033 | 0.0042 | -0.0009 | -0.0005 | 0.0024 | 0.0037 | -0.0012 | -0.0011 |
| 15 | 0.0037 | 0.0039 | -0.0002 | -0.0010 | 0.0023 | 0.0034 | -0.0011 | -0.0008 |
| 16 | 0.0040 | 0.0040 | 0.0000 | -0.0012 | 0.0031 | 0.0033 | -0.0002 | -0.0027 |
| 17 | 0.0040 | 0.0041 | -0.0001 | 0.0003 | 0.0034 | 0.0038 | -0.0004 | -0.0017 |
| 18 | 0.0035 | 0.0039 | -0.0003 | -0.0004 | 0.0025 | 0.0033 | -0.0008 | -0.0007 |
| 19 | 0.0038 | 0.0041 | -0.0003 | -0.0004 | 0.0030 | 0.0037 | -0.0007 | -0.0006 |
| 20 | 0.0038 | 0.0040 | -0.0002 | 0.0005 | 0.0031 | 0.0033 | -0.0002 | -0.0022 |
| 21 | 0.0034 | 0.0038 | -0.0004 | -0.0006 | 0.0030 | 0.0034 | -0.0004 | -0.0010 |
| Sum | 0.0791 | 0.0836<sup>1</sup> | -0.0045<sup>2</sup> | -0.0037<sup>3</sup> | 0.0655 | 0.0736<sup>1</sup> | -0.0081<sup>2</sup> | -0.0348<sup>3</sup> |
| Whole-genome<sup>1+2+3</sup> | 0.0754 |  |  |  | 0.0307 |  |  |  |

**Figure S1:**
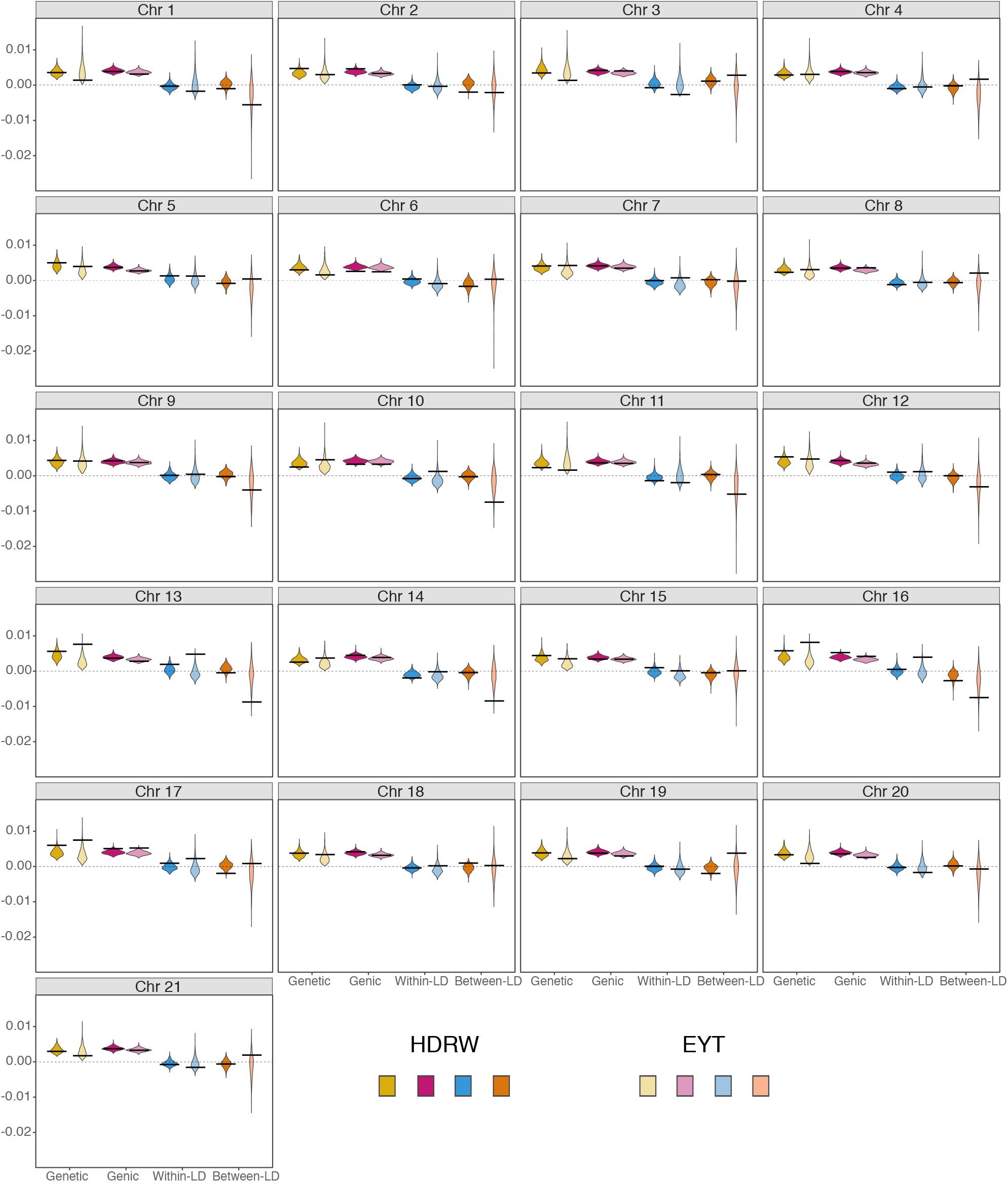
Genetic and genic variances, and within- and between-chromosome linkage dis-equilibrium (LD) covariances by chromosome with the full Bayesian approach for headrows (HDRW, year 18) and elite yield trial (EYT, year 21) (see Fig. 2); black lines denote true values and violins depict posterior distributions

**Figure S2:**
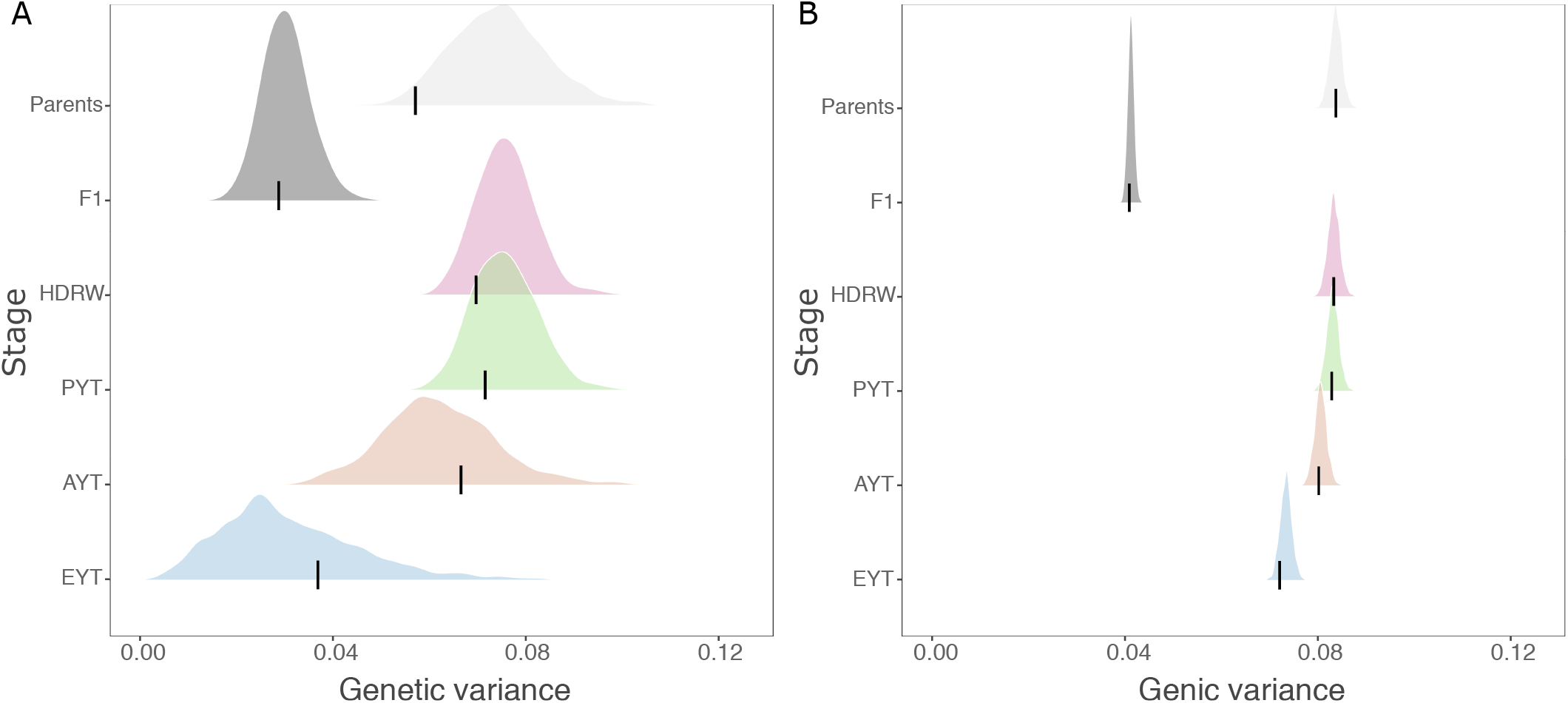
Genetic (A) and genic (B) variance estimated with the empirical Bayesian approach for parents in year 16, F_1_ progeny (F1) in year 17, headrows (HDRW) in year 18, preliminary yield trial (PYT) in year 19, advanced yield trial (AYT) in year 20, and elite yield trial (EYT) in year 21; black lines denote the true values and densities depict posterior distributions

**Figure S3:**
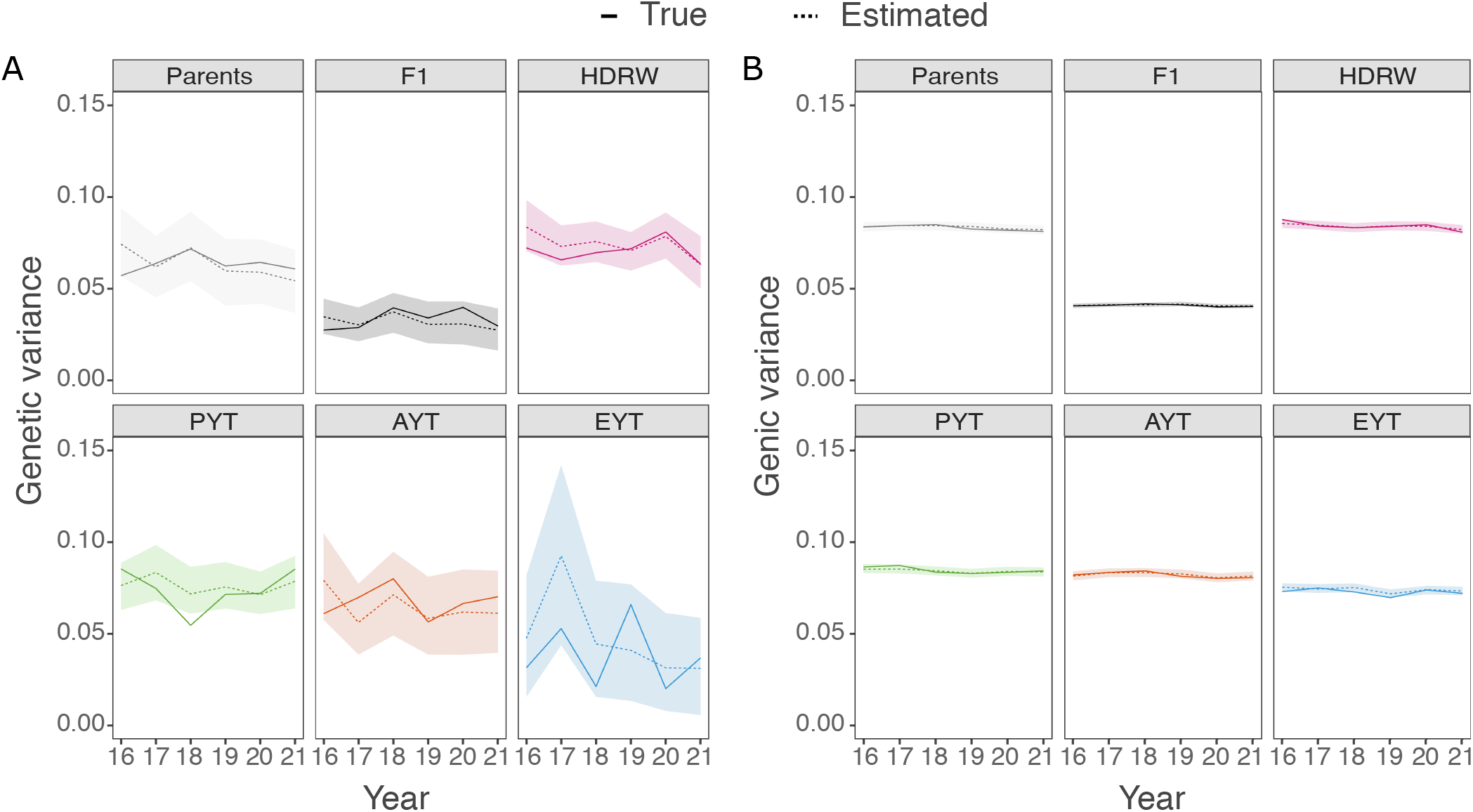
Temporal trend in genetic (A) and genic (B) variance estimated with the empirical Bayesian approach for parents, F_1_ progeny (F1), headrows (HDRW), preliminary yield trial (PYT), advanced yield trial (AYT), and elite yield trial (EYT); solid lines denote the true value, dashed lines denote posterior means and polygons depict 95% posterior quantiles

## References

Akdemir, D., W. Beavis, R. Fritsche-Neto, A. K. Singh, and J. Isidro-Sánchez, 2019 Multi-objective optimized genomic breeding strategies for sustainable food improvement. Heredity 122: 672.

Akdemir, D. and J. I. Sánchez, 2016 Efficient breeding by genomic mating. Frontiers in genetics 7: 210.

Allier, A., S. Teyssèdre, C. Lehermeier, B. Claustres, S. Maltese, et al., 2019 Assessment of breeding programs sustainability: application of phenotypic and genomic indicators to a north european grain maize program. Theoretical and Applied Genetics 132: 1321–1334.

Awata, L. A., P. Tongoona, E. Danquah, B. E. Efie, and P. W. Marchelo-Dragga, 2018 Common mating designs in agricultural research and their reliability in estimation of genetic parameters. IOSR J. Agric. Vet. Sci 11: 16–36.

Bernardo, R., 1994 Prediction of maize single-cross performance using RFLPs and information from related hybrids. Crop Science 34: 20–25.

Bernardo, R., 1996 Best linear unbiased prediction of maize single-cross performance. Crop Science 36: 50–56.

Bernardo, R., 2002 Breeding for quantitative traits in plants, volume 1. Stemma press Woodbury.

Brooks, S., A. Gelman, G. Jones, and X.-L. Meng, 2011 Handbook of Markov Chain Monte Carlo. CRC press.

Bulmer, M., 1971 The stability of equilibria under selection. Heredity 27: 157–162.

Bulmer, M., 1976 The effect of selection on genetic variability: a simulation study. Genetics Research 28: 101–117.

de los Campos, G., J. M. Hickey, R. Pong-Wong, H. D. Daetwyler, and M. P. L. Calus, 2013 Whole-genome regression and prediction methods applied to plant and animal breeding. Genetics 193: 327–345.

de los Campos, G., D. Sorensen, and D. Gianola, 2015 Genomic heritability: what is it? PLoS Genetics 11: e1005048.

Efron, B., 1996 Empirical Bayes methods for combining likelihoods. Journal of the American Statistical Association 91: 538–550.

Falconer, D. S. and T. F. Mackay, 1996 Introduction to quantitative genetics. Longman.

Gaynor, R. C., G. Gorjanc, A. R. Bentley, E. S. Ober, P. Howell, et al., 2017 A two-part strategy for using genomic selection to develop inbred lines. Crop Science 57: 2372–2386.

Gaynor, R. C., G. Gorjanc, and J. M. Hickey, 2020 AlphaSimR: An R-package for breeding program simulations. bioRxiv p. 2020.08.10.245167.

Gianola, D., G. de los Campos, W. G. Hill, E. Manfredi, and R. Fernando, 2009 Additive genetic variability and the Bayesian alphabet. Genetics 183: 347–363.

Gilks, W. R., S. Richardson, and D. Spiegelhalter, 1995 Markov chain Monte Carlo in practice. Chapman and Hall/CRC.

Gorjanc, G., P. Bijma, and J. M. Hickey, 2015 Reliability of pedigree-based and genomic evaluations in selected populations. Genetics Selection Evolution 47: 1–14.

Gorjanc, G., R. C. Gaynor, and J. M. Hickey, 2018 Optimal cross selection for long-term genetic gain in two-part programs with rapid recurrent genomic selection. Theoretical and applied genetics 131: 1953–1966.

Gorjanc, G. and J. M. Hickey, 2019 AlphaBayes: Software for genome-wide marker regression along with fixed and random effects. User Manual. University of Edinburgh, UK.

Gorjanc, G., J. Jenko, S. J. Hearne, and J. M. Hickey, 2016 Initiating maize pre-breeding programs using genomic selection to harness polygenic variation from landrace populations. BMC Genomics 17: 1–15.

Hastie, T. and R. Tibshirani, 2004 Efficient quadratic regularization for expression arrays. Biostatistics 5: 329–340.

Hayes, B. J., P. M. Visscher, and M. E. Goddard, 2009 Increased accuracy of artificial selection by using the realized relationship matrix. Genetics research 91: 47–60.

Hem, I. G., M. L. Selle, G. Gorjanc, G.-A. Fuglstad, and A. Riebler, 2020 Robust genomic modelling using expert knowledge about additive, dominance and epistasis varia-tion. bioRxiv.

Henderson, C. R., 1976 A simple method for computing the inverse of a numerator relationship matrix used in prediction of breeding values. Biometrics pp. 69–83.

Hidalgo, J., S. Tsuruta, D. Lourenco, Y. Masuda, Y. Huang, et al., 2020 Changes in genetic parameters for fitness and growth traits in pigs under genomic selection. Journal of Animal Science 98: skaa032.

Hill, W. G., 2016 Is continued genetic improvement of livestock sustainable? Genetics 202: 877–881.

Hill, W. G., M. E. Goddard, and P. M. Visscher, 2008 Data and theory point to mainly additive genetic variance for complex traits. PLoS Genet 4: e1000008.

Jordan, A., F. Krüger, and S. Lerch, 2019 Evaluating probabilistic forecasts with scoringRules. Journal of Statistical Software 90: 1–37.

Kelleher, J., Y. Wong, A. W. Wohns, C. Fadil, P. K. Albers, et al., 2019 Inferring whole-genome histories in large population datasets. Nature genetics 51: 1330–1338.

Kennedy, B., L. Schaeffer, and D. Sorensen, 1988 Genetic properties of animal models. Journal of Dairy Science 71: 17–26.

Kenward, M. G. and J. H. Roger, 1997 Small sample inference for fixed effects from restricted maximum likelihood. Biometrics 53: 983–997.

Lehermeier, C., G. de Los Campos, V. Wimmer, and C.-C. Schön, 2017 Genomic variance estimates: With or without disequilibrium covariances? Journal of Animal Breeding and Genetics 134: 232–241.

Listgarten, J., C. Lippert, C. M. Kadie, R. I. Davidson, E. Eskin, et al., 2012 Improved linear mixed models for genome-wide association studies. Nature Methods 9: 525–526.

Lush, J., 1937 Animal breeding plans. Iowa State College Press.

Lynch, M. and B. Walsh, 1998 Genetics and analysis of quantitative traits, volume 1. Sinauer Sunderland, MA.

Mather, K. and J. L. Jinks, 2013 Biometrical genetics: The study of continuous variation. Springer.

Meuwissen, T., B. Hayes, and M. Goddard, 2001 Prediction of total genetic value using genome-wide dense marker maps. Genetics 157: 1819–1829.

Meyer, K., 1985 Maximum likelihood estimation of variance components for a multivariate mixed model with equal design matrices. Biometrics pp. 153–165.

Meyer, K. and W. G. Hill, 1997 Estimation of genetic and phenotypic covariance functions for longitudinal or repeated records by restricted maximum likelihood. Livestock Production Science 47: 185–200.

Oakey, H., A. Verbyla, W. Pitchford, B. Cullis, and H. Kuchel, 2006 Joint modeling of additive and non-additive genetic line effects in single field trials. Theoretical and Applied Genetics 113: 809–819.

Oakey, H., A. P. Verbyla, B. R. Cullis, X. Wei, and W. S. Pitchford, 2007 Joint modeling of additive and non-additive (genetic line) effects in multi-environment trials. Theoretical and Applied Genetics 114: 1319–1332.

Ødegård, J., U. Indahl, I. Strandén, and T. H. Meuwissen, 2018 Large-scale genomic prediction using singular value decomposition of the genotype matrix. Genetics Selection Evolution 50: 6.

Peterson, R. E., K. Kuchenbaecker, R. K. Walters, C.-Y. Chen, A. B. Popejoy, et al., 2019 Genome-wide association studies in ancestrally diverse populations: opportunities, methods, pitfalls, and recommendations. Cell 179: 589–603.

Piepho, H. P., J. Möhring, A. E. Melchinger, and A. Büchse, 2008 BLUP for phenotypic selection in plant breeding and variety testing. Euphytica 161: 209–228.

R Core Team, 2019 R: A Language and Environment for Statistical Computing. R Foundation for Statistical Computing, Vienna, Austria.

Schreck, N., H.-P. Piepho, and M. Schlather, 2019 Best prediction of the additive genomic variance in random-effects models. Genetics 213: 379–394.

Selle, M. L., I. Steinsland, J. M. Hickey, and G. G, 2019 Flexible modelling of spatial variation in agricultural field trials with the R package INLA. Theoretical and Applied Genetics 132: 3277–3293.

Smith, A. B., B. R. Cullis, and R. Thompson, 2005 The analysis of crop cultivar breeding and evaluation trials: an overview of current mixed model approaches. The Journal of Agricultural Science 143: 449–462.

Sorensen, D., R. Fernando, and D. Gianola, 2001 Inferring the trajectory of genetic variance in the course of artificial selection. Genetics Research 77: 83–94.

Sorensen, D. and D. Gianola, 2007 Likelihood, Bayesian, and MCMC methods in quantitative genetics. Springer Science & Business Media.

Sorensen, D. and B. Kennedy, 1984 Estimation of genetic variances from unselected and selected populations. Journal of Animal Science 59: 1213–1223.

Strandén, I. and D. J. Garrick, 2009 Technical note: derivation of equivalent computing algorithms for genomic predictions and reliabilities of animal merit. Journal of Dairy Science 92: 2971–2975.

Thompson, R., 2019 Desert island papers a life in variance parameter and quantitative genetic parameter estimation reviewed using 16 papers. Journal of Animal Breeding and Genetics 136: 230–242.

Thompson, R., S. Brotherstone, and I. M. White, 2005 Estimation of quantitative genetic parameters. Philosophical Transactions of the Royal Society B: Biological Sciences 360: 1469–1477.

Tibshirani, R., 1996 Regression shrinkage and selection via the lasso. Journal of the Royal Statistical Society: Series B (Methodological) 58: 267–288.

Tolhurst, D. J., K. L. Mathews, A. B. Smith, and B. R. Cullis, 2019 Genomic selection in multi-environment plant breeding trials using a factor analytic linear mixed model. Journal of Animal Breeding and Genetics 136: 279–300.

Tsuruta, S., I. Misztal, and T. Lawlor, 2004 Genetic correlations among production, body size, udder, and productive life traits over time in holsteins. Journal of dairy science 87: 1457–1468.

Tusell, L., P. Pérez-Rodríguez, S. Forni, X.-L. Wu, and D. Gianola, 2013 Genome-enabled methods for predicting litter size in pigs: a comparison. Animal 7: 1739–1749.

van den Berg, I., I. M. MacLeod, C. M. Reich, E. J. Breen, and J. E. Pryce, 2020 Optimizing genomic prediction for australian red dairy cattle. Journal of Dairy Science 103: 6276–6298.

VanRaden, P. M., 2008 Efficient methods to compute genomic predictions. Journal of dairy science 91: 4414–4423.

Varona, L., A. Legarra, M. A. Toro, and Z. G. Vitezica, 2018 Non-additive effects in genomic selection. Frontiers in Genetics 9: 78.

Walsh, B. and M. Lynch, 2018 Evolution and selection of quantitative traits. OUP Oxford.

Whittaker, J. C., R. Thompson, and M. C. Denham, 2000 Marker-assisted selection using ridge regression. Genetics Research 75: 249–252.

Woolliams, J., P. Berg, B. Dagnachew, and T. Meuwissen, 2015 Genetic contributions and their optimization. Journal of Animal Breeding and Genetics 132: 89–99.

Wright, S., 1931 Evolution in Mendelian populations. Genetics 16: 97.

